# DNA methylation protects cancer cells against senescence

**DOI:** 10.1101/2024.08.23.609297

**Authors:** Xiaoying Chen, Kosuke Yamaguchi, Brianna Rodgers, Delphine Goehrig, David Vindrieux, Xavier Lahaye, Matthieu Nolot, Laure Ferry, Nadine Martin, Pierre Dubus, Fumihito Miura, Takashi Ito, Nicolas Manel, Masato Kanemaki, David Bernard, Pierre-Antoine Defossez

## Abstract

Inhibitors of DNA methylation such as 5-aza-deoxycytidine are widely used in experimental and clinical settings. However, their mechanism of action is such that DNA damage inevitably co-occurs with loss of DNA methylation, making it challenging to discern their respective effects. Here we deconvolute the effects of decreased DNA methylation and DNA damage on cancer cells, by using degron alleles of key DNA methylation regulators. We report that cancer cells with decreased DNA methylation —but no DNA damage— enter cellular senescence, with G1 arrest, SASP expression, and SA-β-gal positivity. This senescence is independent of p53 and Rb, but involves p21, which is cytoplasmic and inhibits apoptosis, and cGAS, playing a STING-independent role in the nucleus. Xenograft experiments show that tumor cells can be made senescent *in vivo* by decreasing DNA methylation. These findings reveal the intrinsic effects of loss of DNA methylation in cancer cells and have practical implications for future therapeutic approaches.

## Introduction

DNA methylation is an abundant modification in mammals (1). In differentiated murine or human cells, about 80% of cytosines in the CpG sequence context are marked with a methyl group on carbon 5, or CpG methylation. CpG methylation is epigenetic as it is transmitted from one cell to its descendants, and it influences genome activity, without changing the sequence itself.

Different cell types have different methylomes, owing to a dynamic equilibrium between the deposition and removal of DNA methylation (2). Removal can occur by active mechanisms (3), or passively, by failure of the DNA methylation maintenance machinery to reproduce the mark following DNA replication.

DNA methylation maintenance involves two key enzymes (4). The first is the DNA methyltransferase DNMT1, which can act on hemimethylated DNA (5). The activity of DNMT1 is regulated by intramolecular inhibition, which limits its activity on unmethylated DNA (6). During and after replication (7, 8), this activity is enabled thanks to another essential actor: UHRF1 (9, 10). This protein interacts with the DNA replication machinery (11, 12) and binds hemimethylated DNA, it can then mono-ubiquitinate histones and the PCNA-associated protein PAF15; these modified proteins, in turn, bind an auto-inhibitory region in DNMT1 and release its activity (13–15).

Functionally, DNA methylation is linked to gene expression. The DNA methylation of gene bodies correlates positively with their expression level. In contrast, high levels of DNA methylation at CpG-rich promoters cause transcriptional repression. In healthy somatic cells, this transcriptional repression by DNA methylation applies to imprinted genes, certain tissue-specific genes including germline genes, and to repeated elements (2, 16, 17).

The pattern of DNA methylation in cancer cells is different from that of WT cells, combining losses of DNA methylation over large domains called “Partially Methylated Regions”, and foci of hypermethylation, in particular over CpG island promoters, including those of repressed Tumor Suppressor Genes (18, 19). These premises provided the rationale for epigenetic therapy in cancer, with the hypothesis that decreasing DNA methylation in cancer cells may normalize their tumor suppressor gene expression and/or exacerbate their DNA methylation anomalies past a tolerable level (20).

One approach to diminish DNA methylation in cancer cells has proven particularly successful and led to the first FDA-approved epigenetic drug for cancer treatment, Vidaza (21). This approach relies on a cytosine analog, 5-aza-deoxy-cytidine (5-aza-dC), being incorporated into replicating DNA. Active DNMT1 reacting with 5-aza-dC leads to the formation of a covalent adduct, which is then removed by the DNA repair machinery. In the process, the DNMT1 molecules are destroyed, leading to passive DNA demethylation.

5-aza-dC, or its precursor 5-aza-C, have been extremely useful molecular probes to examine the phenotypic consequences of decreasing DNA methylation in cancer cells. Depending on cellular context, dose, and duration of treatment, these consequences in most cases are cell differentiation or cell death (21), with senescence reported in a few rare occurrences (22–25). A key phenotype with consequences for cancer biology and treatment is that 5-aza-dC treatment induces the reactivation of repeated elements, producing cytoplasmic nucleic acids that trigger an interferon response (26, 27).

The numerous studies carried out with 5-aza-C and 5-aza-dC have moved the field of cancer epigenetics forward, yet they also suffer from a serious caveat: 5-aza-dC causes loss of DNA methylation but also DNA damage, and one cannot occur without the other. Therefore, it is challenging to deconvolute the effects of DNA demethylation from those of DNA damage, which itself can induce apoptosis or senescence.

One way to disentangle these effects is to perform loss-of-function studies on proteins that maintain DNA methylation, such as DNMT1 or UHRF1, yet existing methods all have limitations. Constitutive genetic knock-out is not applicable to essential genes and may select for adaptation, conditional knock-out is hampered by delayed kinetics, while RNAi and shRNA often fail to achieve total mRNA depletion, and their effects depend on the protein turnover rate. These various shortcomings can be circumvented by using protein degron approaches (28).

We recently generated degron alleles of DNMT1 and UHRF1 in colorectal cancer cells (29). The rapid, complete, and synchronous protein degradation permitted by this approach allowed us to delineate new molecular mechanisms by which UHRF1 maintains DNA methylation homeostasis (29). In the current study, we have used these degron tools to answer a key question in cancer epigenetics: “What are the consequences to a cancer cell when DNA methylation levels decrease in the absence of DNA damage?”.

We report that cells with lowered DNA methylation —but no DNA damage— go into senescence, with the typical attributes of G1 accumulation, enlarged nuclei, a Senescence-Associated Secretory Phenotype (SASP), and positivity for Senescence-Associated Beta-Galactosidase (SA-β-gal). Mechanistically, this senescence is induced more rapidly by inhibiting UHRF1 than DNMT1. It is independent of p53 and the p16/Rb pathways. Instead, it involves cytoplasmic p21, which protects the senescent cells from apoptosis, and cGAS, acting in the nucleus independently of STING. We observed DNA-demethylation-induced senescence in multiple cancer lines originating from different organs, and demonstrated with xenografts that it also occurs *in vivo*.

These findings reveal the intrinsic effects of loss of DNA demethylation in cancer cells, and have practical implications for future therapeutic approaches.

## Results

### Prolonged depletion of DNMT1 and/or UHRF1 triggers senescence in colorectal cancer cell lines

To examine the consequences of prolonged DNA demethylation, we used the colorectal cancer cell line HCT116, exploiting the auxin-inducible degron (AID1) system (30). As described in our previous publication (29), these HCT116 cells (WT, U^AID1^, D^AID1^ and UD^AID1^) express the plant E3 ligase OsTIR1 under a doxycycline (Dox)-inducible promoter, and were edited so that UHRF1, DNMT1, or both, had a mini-AID (mAID) tag, allowing for the proteins of interest to be rapidly degraded in the presence of auxin (Figure 1A). A time-course western blot proved that UHRF1 and/or DNMT1 were completely degraded throughout the duration of the experiment (Figure 1B). To characterize the effect of chronic depletion of UHRF1, DNMT1, or both, we treated the HCT116 WT and its derivatives with Dox/Auxin continuously for 8 days and quantified cell proliferation. The cells depleted of DNMT1 displayed a significant growth defect compared to the WT; this defect was worse in UHRF1-depleted cells, and worst in cells lacking both DNMT1 and UHRF1 (Figure 1C).

**Figure 1.**
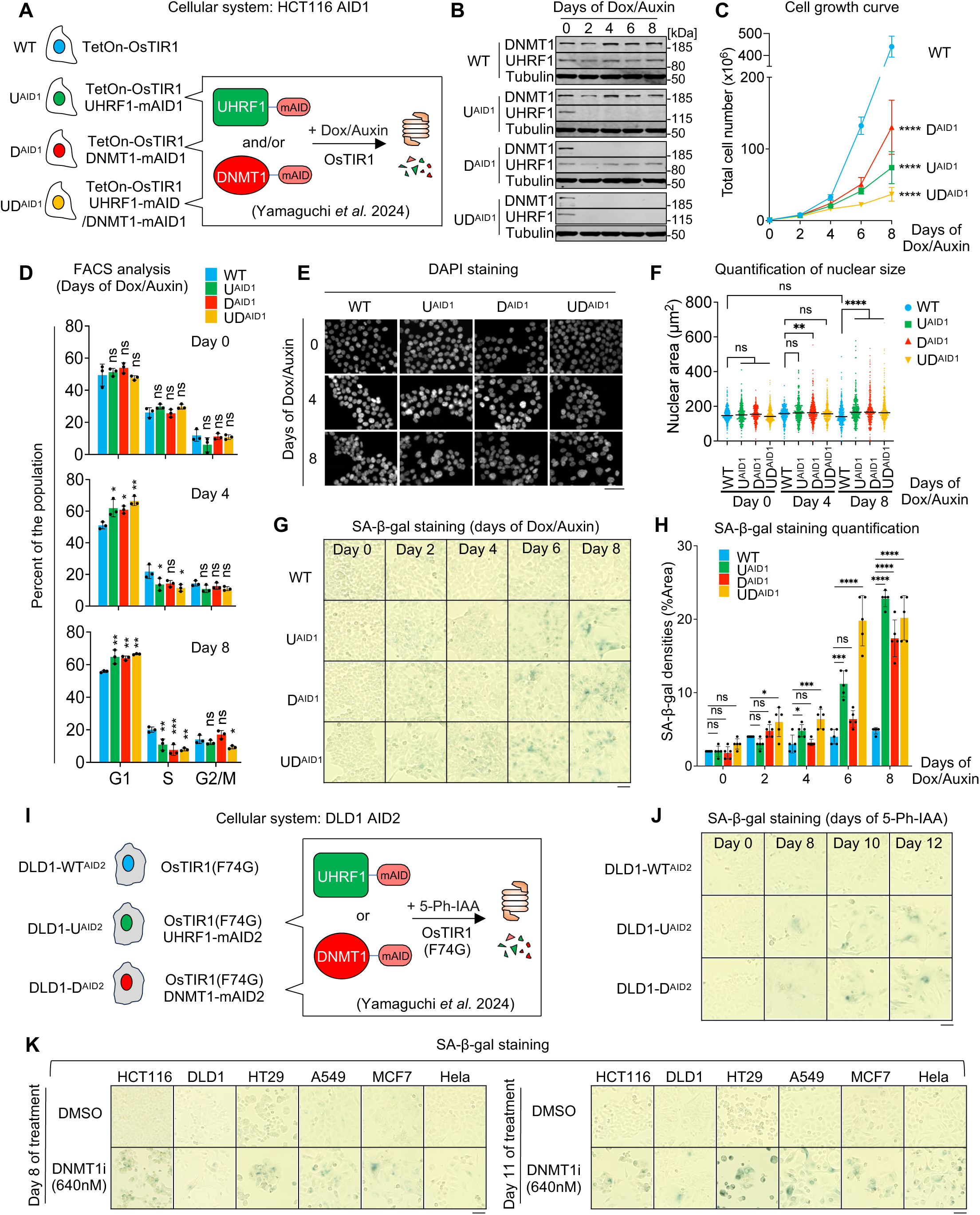
Prolonged UHRF1 or DNMT1 depletion triggers senescence in cancer cells. (A) The auxin degron system in HCT116 cells conditionally expressing OsTIR1. (B) Western blot validating the degradation of UHRF1 and/or DNMT1 upon Dox/Auxin treatment. (C) Cell growth curve, N = 3 biological replicates. (D) Cell-cycle analysis by FACS with BrdU and PI staining. N = 3 biological replicates. (E) Representative images of DAPI staining in the indicated lines. (F) Nuclear area determination by DAPI staining. (G) Representative images of SA-β-gal staining after Dox/Auxin treatment. (H) Quantification of SA-β-gal staining. N = 5 fields of view. (I) The auxin degron system in DLD1 cells constitutively expressing OsTIR1 (F74G). (J) Representative images of SA-β-gal staining after 5-Ph-IAA (1 µM) treatment. (K) Representative images of SA-β-gal staining after treatment of the indicated cells with DMSO or DNMT1 inhibitor GSK-3685032 (640 nM) for the indicated duration. All scale bars are 50 µm. Data of (C), (D), and (H) are presented as mean ± SD and analyzed by one-way ANOVA test with Dunnett’s multiple comparisons test. Data of (F) are presented as median and analyzed by Kruskal-Wallis test with Dunn’s multiple comparisons test. In all figures we use the following convention: * p < 0.05, ** p < 0.01, *** p < 0.001, **** p < 0.0001, ns: non-significant.

To further explore this proliferation defect, we next performed a cell-cycle analysis. HCT116 WT, U^AID1^, D^AID1^ and UD^AID1^ cells were collected at 0, 4, or 8 days of Dox/Auxin treatment, and stained with bromodeoxyuridine (BrdU) and propidium iodide (PI) to profile the cell-cycle proportions. Through analysis with flow cytometry, we observed a significant decrease in the population of cells in the S phase of the cell cycle after UHRF1 or DNMT1 depletion compared to the WT (Figure 1D), which parallels the loss of proliferation seen in the cell growth curve. We also noted a significant accumulation of the cell population in the G1 phase after UHRF1 and/or DNMT1 removal (Figure 1D).

The reduced proliferation of cells lacking UHRF1 and/or DNMT1 could involve increased apoptosis and/or increased senescence, so we went on to examine both phenotypes. We began by performing an Annexin V analysis, in which the cells are categorized into live, early apoptotic, or late apoptotic/necrotic profiles through analysis with flow cytometry (Figure S1A). When comparing day 0 to day 8 of auxin treatment, we detected no extensive increase in apoptosis (Figure S1B). For further confidence in excluding apoptosis, we performed a TUNEL assay followed by fluorescent microscopy to observe potential DNA fragmentation as seen in apoptotic cells. After 8 days of auxin treatment, the cells depleted for UHRF1 and/or DNMT1 did not display any detectable staining for fragmented DNA (Figure S1C), unlike the positive control (cells treated with DNase) (Figure S1D). These data support the notion that the loss of cell growth upon UHRF1 and/or DNMT1 depletion is not caused by increased cell death, leaving increased senescence as another possible explanation.

Senescent cells display a number of morphological alterations, including enlarged nuclei (31). After quantification of the DAPI signal in the various experimental conditions, we determined that indeed the loss of UHRF1 and DNMT1 resulted in increased nuclear size (Figures 1E, 1F), further suggesting a senescent phenotype in the degron cells.

To further confirm this possible phenotype, we next assayed senescence-associated-beta-galactosidase (SA-β-gal), a common marker of the increased lysosomal activity present in senescent cells (32). This revealed a significant increase of SA-β-gal positive cells after the depletion of DNMT1, which became more pronounced with time. The increase was stronger in cells lacking UHRF1, and strongest in cells lacking both DNMT1 and UHRF1 (Figures 1G, 1H), which parallels their degree of growth impairment.

These experiments show that HCT116 cells lacking DNMT1, UHRF1, or both, present 3 phenotypes found in senescent cells: accumulation in G1 phase of the cell cycle; enlargement of the nucleus; and expression of SA-β-gal. These data suggest that senescence contributes to the reduced proliferation of cells lacking UHRF1, DNMT1, or both.

### Senescence in the depleted cells is linked to loss of DNA methylation and occurs in multiple cellular backgrounds

The loss of either DNMT1 or UHRF1 leads to senescence in HCT116 cells, suggesting that this response may be caused by decreased DNA methylation. To test this possibility, we rescued the degron cells with previously characterized UHRF1 and DNMT1 variants (29), which do, or do not, support DNA methylation (Figures S1E, S1F, S1H, S1I). We then performed SA-β-gal staining in the rescued cells after 8 days of auxin treatment. We observed a strong correlation between loss of DNA methylation and senescence: the cells in which DNA methylation was sustained (WT or TTD mutant UHRF1, WT or PBD mutant DNMT1) were negative for SA-β-gal staining, while the cells in which DNA methylation was decreased (empty vectors, UBL or PHD or SRA or RING mutant UHRF1, UIM or H3 binding or catalytic mutant DNMT1) stained positively for SA-β-gal staining (Figures S1F, S1G, S1I, S1J). These data suggest that lack of DNA methylation underpins the senescence of HCT116 degron cells in our study.

Next, we sought to determine whether the observations made in HCT116 could be extended to other cellular backgrounds. In our previous publication (29), we generated auxin-inducible degron alleles of UHRF1 and DNMT1 in colorectal DLD1 cells (Figure 1I), using the AID2 system in which the OsTIR1 has the F74G modification and the activator is 5-Ph-IAA (33). We treated the DLD1 cells with 5-Ph-IAA for 8 days and even longer, then performed the staining: this revealed an increase of SA-β-gal positivity after the removal of either UHRF1 or DNMT1 (Figure 1J), extending the link between DNA demethylation and senescence beyond HCT116 cells. We next asked whether the senescence phenotype seen with the removal of UHRF1 or DNMT1 was specific to the degron system, or if we could confirm it with an independent intervention. For this we treated various cancer cell lines (HCT116, DLD1, HT29, A549, MCF7 and HeLa) with a DNMT1-selective noncovalent inhibitor that does not cause DNA damage, GSK3685032 (34). We incubated the cells with 640nM inhibitor for 8 days and 11 days, then performed the SA-β-gal assay (Figure 1K). This inhibition of DNMT1 led to an increase of SA-β-gal positive cells in all cell lines, further validating the hypothesis that DNA demethylation induced by UHRF1 or DNMT1 removal triggers senescence, a phenomenon consistent in multiple cancer cell types as well as with various methods of depletion.

### Transcriptional profiling of degron cells reveals an interferon response and a SASP signature

Having identified this senescence induced by decreased DNA methylation, we moved on to a detailed molecular characterization by performing an RNA-seq timed series (0, 2, 4, 6, and 8 days of Dox/Auxin treatment, Figure 2A). These experiments precise and extend the limited transcriptome analysis we previously reported at days 0 and 4 (29), yielding important new insight as reported below.

**Figure 2.**
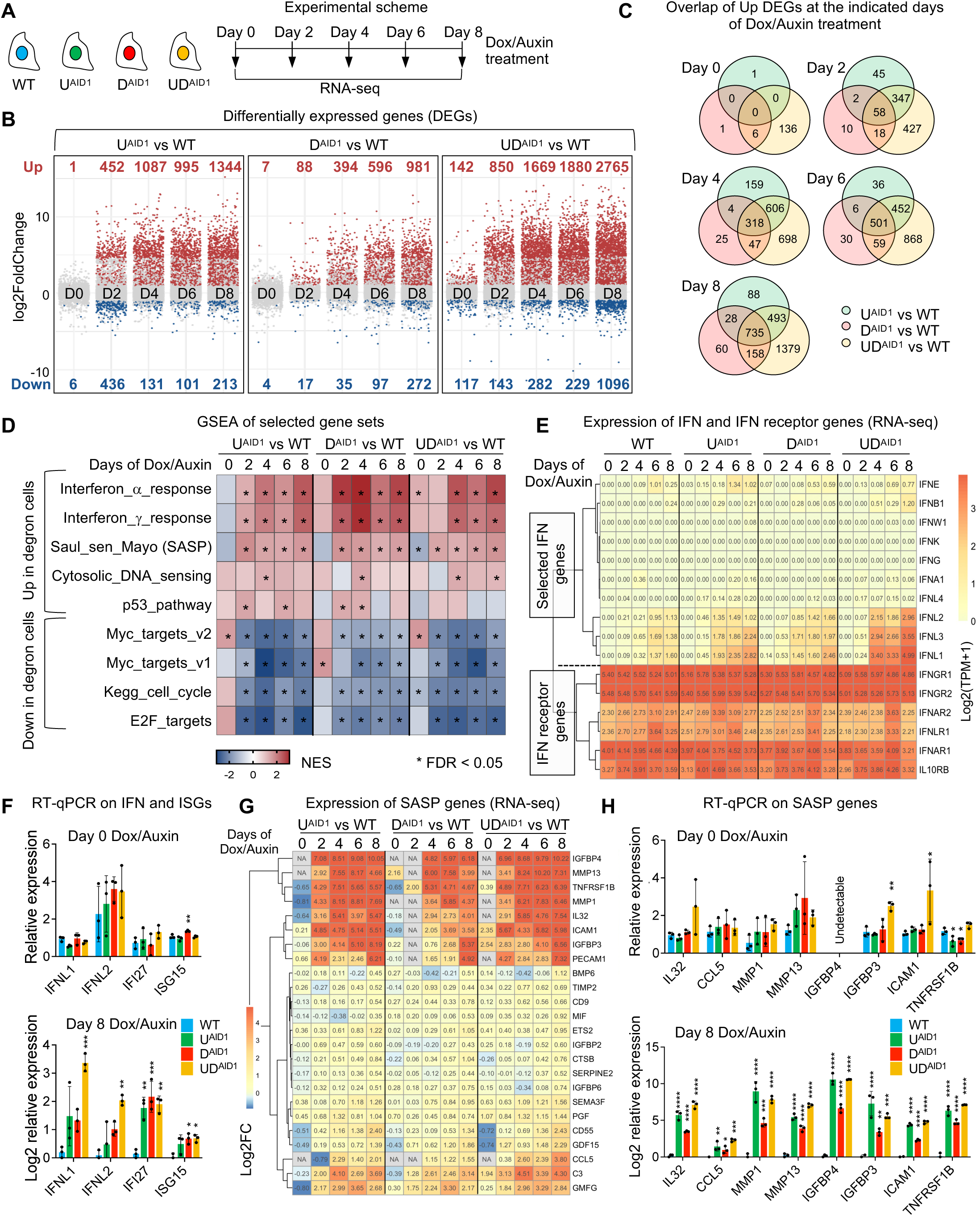
Senescence caused by loss of DNA methylation is accompanied by an interferon response and a SASP signature. (A) Experimental scheme of RNA sequencing. (B) Number of differentially expressed genes (DEGs) at the indicated days of Dox/Auxin treatment. Up-regulated genes (red): adjusted p value < 0.05 and log2 FoldChange > 1. Down-regulated genes (blue): adjusted p value < 0.05 and log2 FoldChange < −1. Light grey dots: no significant change. (C) Venn diagram of significantly up-regulated DEGs. (D) Gene set enrichment analysis (GSEA) of selected gene sets from the Molecular Signatures Database (MSigDB). The heatmap represents the normalized enrichment score (NES) in each comparison at the indicated days. Significantly enriched gene sets are shown in red; significantly depleted gene sets are displayed in blue. Conditions with an asterisk are significant with a false discovery rate (FDR) < 0.05. (E) mRNA abundance for selected interferon (IFN) and IFN receptor genes (using log2-transformed transcripts per million (TPM)). Rows are grouped by hierarchical clustering. (F) RT-qPCR on selected IFN and IFN-stimulated genes (ISGs). N = 3 biological replicates. (G) Changes in mRNA abundance of SASP genes (log2 FoldChange) that are significantly induced (adjusted p value < 0.05) on Day 8 for all comparisons. Rows are grouped by hierarchical clustering. (H) RT-qPCR on selected SASP genes. N = 3 biological replicates. Data of (F) and (H) are presented as mean ± SD and analyzed by one-way ANOVA test with Dunnett’s multiple comparisons test. We use the following convention: * p < 0.05, ** p < 0.01, *** p < 0.001, **** p < 0.0001.

Principal component analysis plots showed that biological triplicates of each condition were highly similar (Figure S2A). Untreated WT and degron lines also display very comparable RNA-seq patterns, meaning that the AID tag has no obvious effect on cells, as we previously reported (29). Finally, the transcriptome patterns of 6-day treated WT and 8-day treated WT cells differed from untreated, 2-day or 4-day treated WT (Figure S2A), reflecting the known effects of auxin treatment, even on WT cells (33). To remove the influence of auxin, in all subsequent analyses, we compared the Dox/Auxin-treated degron lines to the WT counterpart that had been Dox/Auxin-treated for the same duration.

We first computed differentially expressed genes (DEGs) upon UHRF1 and/or DNMT1 depletion over time (Figure 2B). For all degron cells, the number of DEGs increased steadily with time, and there were more upregulated than downregulated genes. The DNMT1 degron cells showed the lowest number of DEGs at all time points, while UHRF1 degron cells had a higher number of DEGs, and the UHRF1/DNMT1 degron cells had the highest number, which parallels both the degree of DNA methylation loss (29) and the prevalence of senescent cells (see previous section). There is a high overlap of genes induced by removing UHRF1, DNMT1, or both (Figure 2C), consistent with the possibility that they are responding to the loss of DNA methylation caused by removing either protein.

We next performed gene set enrichment analysis (GSEA) on our data; a non-exhaustive list of gene sets is shown in Figure 2D. Several signatures were downregulated after Dox/Auxin treatment: those are generally related to cell cycle (E2F targets, Myc targets, cell cycle), which agrees with the decreased cellular proliferation we observed upon treatment of the degron cells. Conversely, many signatures were upregulated in treated cells, and among them were interferon (IFN) response signatures. This aligns with other studies where the interferon response is induced when DNA methylation decreases, after treatment with the DNA demethylating agent such as 5-aza-C or 5-aza-dC (26, 27), or following RNA interference on DNMT1 or UHRF1 (35, 36).

We then sought to more accurately identify the molecular actors involved in the interferon response by a thorough examination of the RNA-seq dataset. For this, we generated a heatmap of mRNA expression for interferon- and interferon receptor-encoding genes (Figure 2E). The genes that encode IFN alpha, beta, gamma, kappa, and omega were never expressed at appreciable levels, while the gene for IFN epsilon (IFNE) showed a modest induction at the latest time points. In contrast, the mRNAs encoding IFN lambda (IFNL1, IFNL2, IFNL3) were induced from day 4 of the time course (Figure 2E). In parallel, we observed that the cells express the genes encoding receptors for various IFN types, including receptors for IFN lambda (IFNLR1 and IL10RB, Figure 2E), and that these levels remained relatively stable during the experiment. To validate these results, we performed RT-qPCR on degron cells at days 0 and 8 of treatment. This experiment confirmed the increase of IFNL1 and IFNL2 (Figure 2F), and demonstrated the induction of IFI27 and ISG15, two genes stimulated by interferon. Furthermore, we collected serum-free conditioned medium (CM) from 8-day Dox/Auxin-treated WT/AID lines and, for a positive control, from PolyI:C transfected cells (Figure S2B). This serum-free CM was utilized in ELISA experiments, which revealed that the treated degron HCT116 cells secrete IFN lambda (Figure S2C).

Upon chronic depletion of UHRF1 and/or DNMT1, the degron cells express both interferon lambda and its receptors, so we hypothesized that they might have paracrine effects and stimulate a secondary interferon response. To test this idea, we supplemented the CM from 8-day auxin treated WT/AID lines, and from cells made senescent by the DNA-damaging agent etoposide with 10% FBS (Figure S2B). Subsequently, we cultured HCT116 WT cells with this CM for 24 hours, extracted total RNA from these cells and performed RT-qPCR analysis. We found that conditioned medium from cells depleted of UHRF1 and/or DNMT1 induced interferon-responsive genes (IRF7, IFI27, ISG15, OAS1, OAS1L) in WT cells, whereas conditioned medium from etoposide-treated cells did not (Figure S1D). In parallel, we have confirmed that the etoposide-treated cells did not have increased IFN lambda gene expression (data not shown). These data further substantiate the interferon response of the degron cells.

Our GSEA analysis (Figure 2D) used a SASP signature containing 125 genes (37). We refined the SASP signature specific to our system by examining the expression of individual SASP genes in the RNA-seq data (Figure 2G). All three degron lines displayed a similar SASP pattern: they induced IL32 and other cytokines, CCL5 and other chemokines, MMP1, MMP13, and other proteases, IGFBP3, IGFBP4 and other growth factors, and ICAM1, TNFRSF1B, and other receptors. These findings from RNA-seq analysis were confirmed by RT-qPCR (Figure 2H). This SASP signature was induced fastest and strongest in the UD^AID1^ degron cells, more slowly and less strongly in the U^AID1^ degron cells, and was more delayed and subdued in the D^AID1^ degron cells (Figure 2G), which again parallels the kinetics of DNA demethylation and senescent phenotypes. The SASP phenotype is not restricted to HCT116 degron cells, as we also observed the induction of the same SASP genes in DLD1 degron cells upon prolonged UHRF1 or DNMT1 depletion (Figure S2E).

Collectively, our transcriptome data show that the loss of DNA methylation caused by removal of DNMT1, UHRF1, or both, causes HCT116 cells to display an ISG response and a SASP signature. This signature is accompanied by activation of different signaling pathways, including p53 (Figure 2D), therefore we next investigated the determinants of SASP activation and senescence appearance.

### DNA demethylation-induced senescence is not accompanied by DNA damage and is independent of p53 and p16/Rb

One of the established triggers of cellular senescence is DNA damage, which can act by activating p53 (38). To understand whether the senescence observed in the UHRF1/DNMT1-depleted HCT116 cells may involve DNA damage, we first carried out immunofluorescence on degron cells with an antibody for γ-H2AX, a marker of DNA double-strand breaks (Figure 3A). As a positive control, we treated cells with hydroxyurea (HU), which interferes with DNA replication; this gave rise to a clear γ-H2AX signal, as expected. In contrast, even upon prolonged Dox/Auxin treatment (8 days), the loss of UHRF1 and/or DNMT1 did not lead to any detectable increase of the γ-H2AX signal (Figures 3A, S3B). The γ-H2AX signal examined by immunofluorescence in cells at the intermediate 4-day timepoint was also below the detection threshold (Figures S3A, S3B). To confirm these data, we used western blotting. Two positive controls were used: treatment with HU, or with etoposide, which inhibits topoisomerase II thus causing double-strand breaks. We tested 4 different actors of the DNA damage response: γ-H2AX, Chk1, Chk2, and p53. All four proteins behaved as anticipated in the positive controls, with increases of γ-H2AX, phospho-Chk1, phospho-Chk2, and p53 (Figure 3B). In contrast, none of these four markers were induced in the auxin-treated degron cells relative to the auxin-treated WT cells (Figure 3B). Altogether these data suggest that DNA damage is unlikely to be a major driver of senescence in the degron cells.

**Figure 3.**
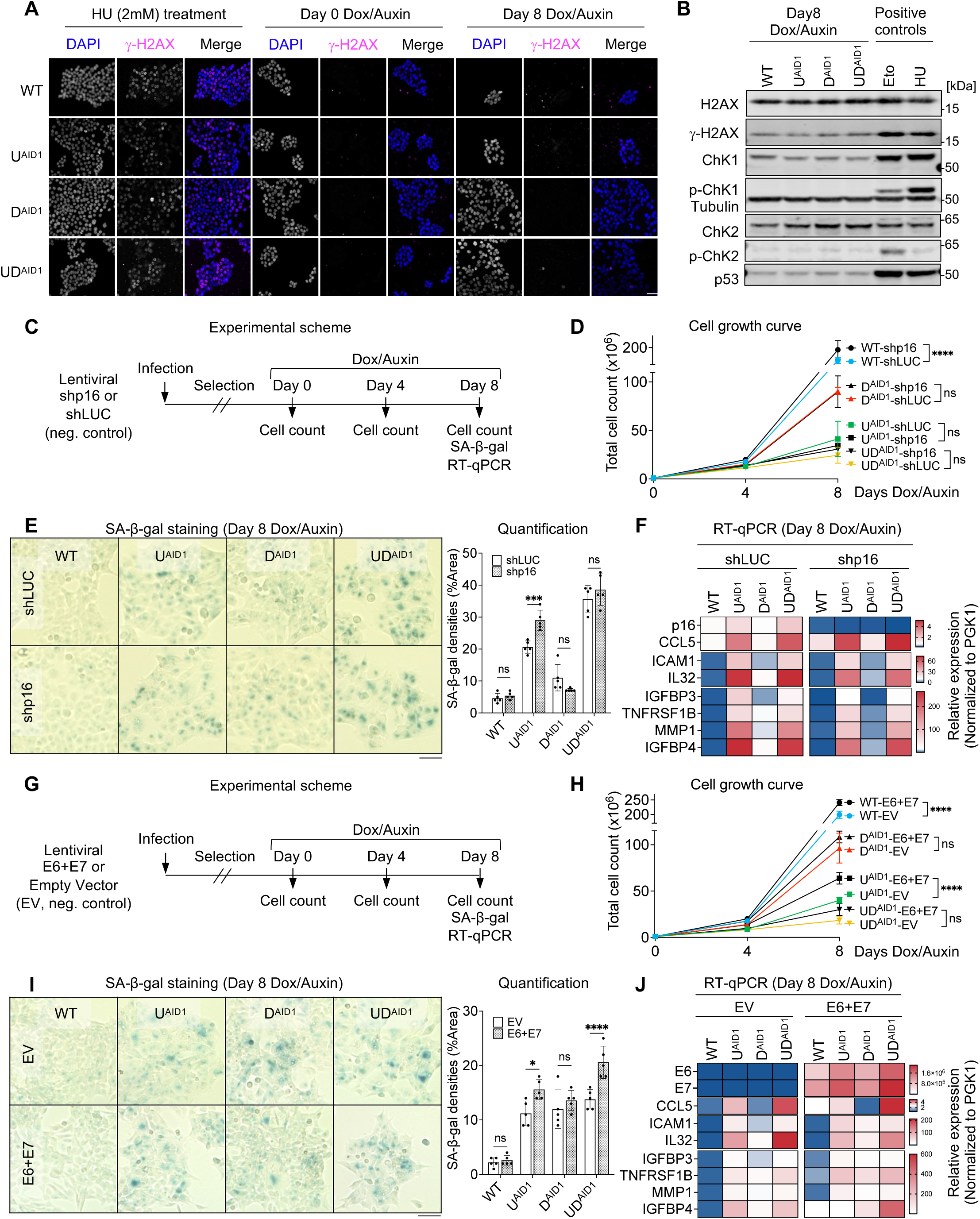
Senescence caused by loss of DNA methylation is independent of p53 and p16/Rb. (A) γ-H2AX Immunofluorescence in HCT116 lines upon hydroxyurea treatment (HU, 2 mM, 4 hours) or upon Dox/Auxin treatment. (B) Immunoblots for total H2AX, γ-H2AX, Chk1, phosphorylated Chk1 (Ser317), Chk2, phosphorylated Chk2 (Thr68), and p53 in HCT116 lines after 8-day Dox/Auxin treatment. HCT116 WT cells treated with etoposide (Eto, 10 µM, 4 hours) or hydroxyurea (HU, 2 mM, 4 hours) used as positive controls. (C) Scheme of p16 depletion experiments. (D) Total cell numbers at the indicated days of Dox/Auxin treatment. N = 3 technical replicates. (E) SA-β-gal staining and quantification of indicated lines. N = 5 fields of view. (F) RT-qPCR results of p16 (CDKN2A) and selected SASP genes in indicated conditions. (G) Scheme of E6+E7 expression experiments. (H) Total cell numbers at the indicated days of Dox/Auxin treatment. N = 3 technical replicates. (I) SA-β-gal staining and quantification of indicated lines. N = 5 fields of view. (J) RT-qPCR results of E6, E7 and selected SASP genes in indicated conditions. All scale bars are 50 µm. Data of (D), (E), (H) and (I) are presented as mean ± SD and analyzed by two-way ANOVA with Sidak’s multiple comparisons test. Heatmap data of (F) and (J) are presented as mean from 3 technical replicates. In all figures we use the following convention: * p < 0.05, *** p < 0.001, **** p < 0.0001, ns: non-significant.

To further delineate the senescence mechanisms active in our cells, we next investigated the p16/Rb pathway. p16/INK4a inhibits Cyclin D/CDK complexes, thus preventing the phosphorylation of Rb and entry into S phase, and the activation of the p16/Rb pathway is frequently observed in senescent cells (39). We first analyzed the expression of CDKN2A gene, which encodes p16 in HCT116 cells depleted for UHRF1 and/orDNMT1 using RT-qPCR, and found that CDKN2A was significantly upregulated in the U^AID1^ and UD^AID1^ cells after 8 days of Dox/Auxin treatment (Figure S3C). Western blotting confirmed that p16 was induced at the protein level in these samples (Figure S3D). In parallel, we inspected Rb expression and phosphorylation, and found both Rb and phospho-Rb protein levels decreased upon the removal of either UHRF1 or UHRF1 and DNMT1 (Figure S3E). This suggests that the p16/Rb pathway may be involved in driving cell cycle arrest and senescence in our system.

To test this possibility, we applied an shRNA approach to deplete p16. The WT and degron cells were infected with a lentiviral vector expressing shp16, or the negative control shLuc, then selected to ensure permanent shRNA expression. After these steps, the cells were treated with Dox/Auxin and we assessed their growth, SA-β-gal activity, and gene expression pattern by RT-qPCR (Figure 3C). The knockdown of p16 did not ameliorate the slow proliferation of degron cells (Figure 3D), nor their increased SA-β-gal positivity (Figure 3E). RT-qPCR confirmed sustained suppression of the p16 gene following an 8-day Dox/Auxin treatment, yet the knockdown did not mitigate SASP gene expression (Figure 3F). Collectively, these results indicate that p16 is not a key enforcer of cell cycle arrest or senescence in cells lacking UHRF1 and/or DNMT1.

To further test the contribution of p53 and Rb to senescence in our system, we exploited the oncogenic proteins of human papilloma virus HPV16: E6, that degrades p53, and E7, that inactivates Rb. The WT and degron cells were infected with an empty lentiviral vector, or with a vector expressing E6 and E7, and selected to ensure stable expression (Figure 3G). We verified that the p53 protein was degraded upon E6 and E7 expression (Figure S3F). As expected, the inactivation of p53 and Rb caused WT cells to grow faster (Figure 3H); in contrast this treatment did not rescue the proliferation defect of D^AID1^ or UD^AID1^ lines treated with auxin (Figure 3H). The treatment ameliorated the growth of U^AID1^ cells treated with auxin, yet they remained severely growth-impaired (Figure 3H). Moreover, the inactivation of p53 and Rb failed to reduce the SA-β-gal positivity (Figure 3I) or SASP gene expression in the treated degron cells (Figure 3J).

Taken together, these results establish that the senescence observed in HCT116 after prolonged DNA demethylation is independent of the p53 and p16/Rb pathways. This fits with the observations of Figure 1K, showing senescence induced by loss of DNA methylation in cells lacking functional p53 (DLD1, HT29, HeLa) and/or Rb (HeLa).

### p21 is found in the cytoplasm of cells with decreased DNA methylation and contributes to their resistance against apoptosis

Having excluded the role of canonical pathways (p53, p16/Rb) in the senescence of cells chronically deprived of UHRF1 and/or DNMT1, we investigated the potential role of other factors. We first exploited our RNA-seq data and observed that similarly to CDKN2A/p16, CDKN1A/p21 was also elevated at various time points in all the degron lines, and we confirmed this observation by RT-qPCR (Figure 4A). p21 was also induced in the degron lines after 8-day Dox/Auxin treatment at protein level (Figure 4B), indicating a potential role in the senescence process.

**Figure 4.**
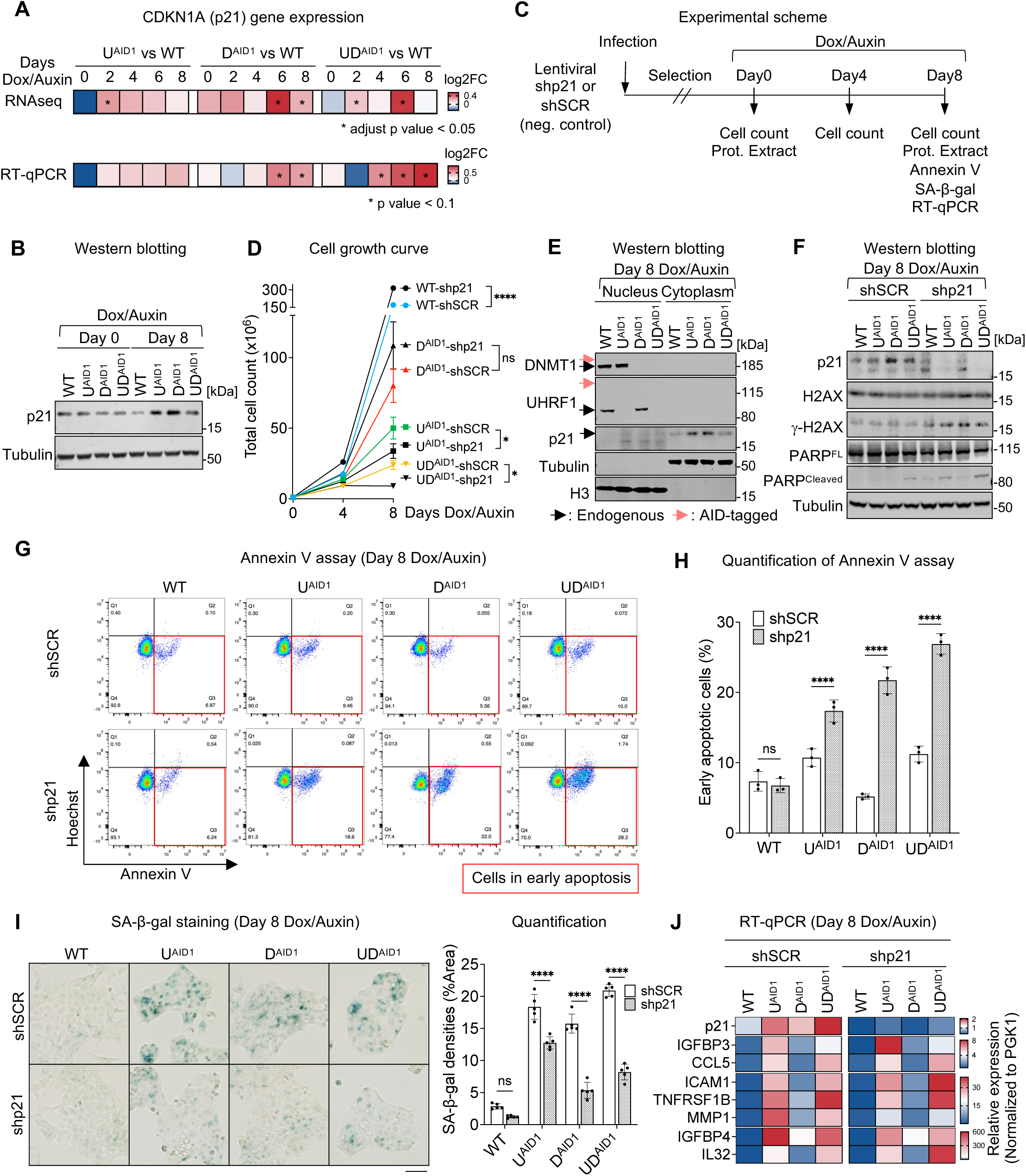
p21 contributes to resistance against apoptosis during senescence caused by decreased DNA methylation. (A) Time-course analysis of CDKN1A (p21) mRNA abundance, presented as log2 fold change in RNA-seq data (upper panel) and RT-qPCR results (lower panel). (B) Immunoblots for p21 in the indicated condition. (C) Experimental scheme of p21 depletion and assessment. (D) Total cell numbers at the indicated days of Dox/Auxin. N = 3 technical replicates. (E) Immunoblots for DNMT1, UHRF1, p21 in the nuclear and cytoplasmic fractions of indicated lines at Day 8 of Dox/Auxin treatment. Black arrows indicate the endogenous proteins of interest and red arrows indicate the endogenous AID-tagged proteins of interest. (F) Immunoblots of p21, H2AX, γ-H2AX, full-length (FL) PARP and cleaved PARP in the indicated lines at Day 8 of Dox/Auxin treatment. (G) Representative results of Annexin V/Hoechst 33342 double staining in the indicated conditions. Cells in early apoptosis are indicated by the red square. (H) Percentages of early apoptotic cells (Annexin V positive/Hoechst 33342 negative). N = 3 technical replicates. (I) SA-β-gal staining (left panel) and quantification (right panel) of indicated lines. N = 5 fields of view. Scale bar is 50 µm. (J) RT-qPCR results of p21 and selected SASP genes in indicated conditions. Data of (D), (H) and (I) are presented as mean ± SD and analyzed by two-way ANOVA with Sidak’s multiple comparisons test. Heatmap data of (J) are presented as mean from 3 technical replicates. We use the following convention: * p < 0.05, **** p < 0.0001, ns: non-significant.

To test this possible role, we generated a lentiviral construct expressing an shRNA targeting p21, stably expressed it in degron cells, then quantified growth and markers of senescence (Figure 4C). We first monitored cell proliferation after p21 depletion and found that the absence of p21 significantly increased the cell growth of WT cells, as expected, but that it decreased the numbers of U^AID1^ and UD^AID1^ lines (Figure 4D), indicative of slower proliferation and/or increased apoptosis in these cells.

CDKN1A/p21 has diverse functions depending on its cellular localization (40): nuclear p21 inhibits various Cyclin/CDK complexes and restricts cell cycle progression, whereas cytoplasmic p21 has pro-survival effects by restraining JNK signaling and caspase activity (41). Cell cycle exit and resistance to apoptosis are both common phenotypes of senescent cells (42), therefore we next assessed the subcellular location of p21 in the degron cells. For this, we separated the nuclear and cytoplasmic fractions of HCT116 cells after 0 and 8-day Dox/Auxin treatment and performed western blotting. Upon validating the depletion of UHRF1 and DNMT1, the results revealed that the majority of the p21 protein present in senescent cells resided in the cytoplasm (Figure 4E, S4A). We then measured the apoptosis marker and DNA damage marker after p21 removal in degron cells, and observed increased abundance of cleaved poly ADP-Ribose Polymerase (PARP), as well as increased γ-H2AX after 8-day Dox/Auxin treatment (Figure 4F), which was not observed in the same conditions at 0-day Dox/Auxin treatment (Figure S4B). Both results are compatible with an anti-apoptotic role of p21 in the cells rendered senescent by lack of DNMT1 and/or UHRF1. To further ascertain this possible role, we directly measured the level of apoptosis in our cells by an Annexin V assay (Figure 4G). Compared to the control conditions, the loss of p21 led to an increase of apoptosis in the conditions depleted for UHRF1 and/or DNMT1 (Figure 4H). These data again support the notion that the senescent cells resist apoptosis in part via p21.

Additionally, p21 knockdown led to a partial reduction of SA-β-gal positivity (Figure 4I), but RT-qPCR showed that p21 knockdown did not decrease SASP expression (Figure 4J).

Collectively, our data show that p21 is located in the cytoplasm of senescing cells with lowered DNA methylation and promotes their survival, but is not involved in the SASP induction. Lastly, we investigated the possible causes of p21 upregulation in cells depleted of UHRF1 and/or DNMT1. The first actor we considered was c-Myc, which is known to repress p21 transcription (43–45). Three lines of evidence suggest that c-Myc activity decreases in the degron cells as they become senescent upon exposure to Dox/Auxin: first, the c-Myc signatures are downregulated in the GSEA analysis (see Figure 2D above); second, the c-Myc activity —as reflected in the VIPER (Virtual Inference of Protein-activity by Enriched Regulon analysis) score (46)— also decreases (Figure S4C); third, the levels of c-Myc protein (Figure S4D) diminish. Therefore, the decrease of c-Myc may account, at least in part, for the induction of p21. Another negative regulator of p21 transcription is TFAP4 (Transcription Factor AP4) (47, 48). Similarly to c-Myc, we saw that the VIPER score of TFAP4 decreases in the degron cells treated with auxin (Figure S4C), concomitant with a decrease abundance of TFAP4 protein (Figure S4D). TFAP4 has been shown in other contexts to prevent senescence (49), so it may play this role in our system as well.

### Nuclear cGAS contributes to the SASP response and lysosomal activity of senescent cells

The nuclear acid sensor pathway cGAS/STING, comprised of cyclic GMP-AMP synthase (cGAS), stimulator of interferon genes (STING), and downstream signaling adaptors, can be involved in cellular senescence and SASP-derived inflammation (50–52), prompting us to examine its possible implication in our system.

To this end, we first performed RT-qPCR on the HCT116 cells at various timepoints after Dox/Auxin treatment to measure the expression of cGAS. We observed that the loss of UHRF1 and DNMT1 together triggered a dramatic upregulation of cGAS; this was less prominent after UHRF1 depletion and not significant after DNMT1 loss (Figure 5A). We further validated this with western blot (Figure 5B), and found that the cGAS protein was undetectable in untreated HCT116 WT cells, in agreement with previous work (53). In contrast, the depletion of UHRF1 and DNMT1, or to a lesser extent the depletion of UHRF1 alone, led to an induction of cGAS protein that was visible as early as day 2 of treatment, and increased continuously over time (Figure 5B).

**Figure 5.**
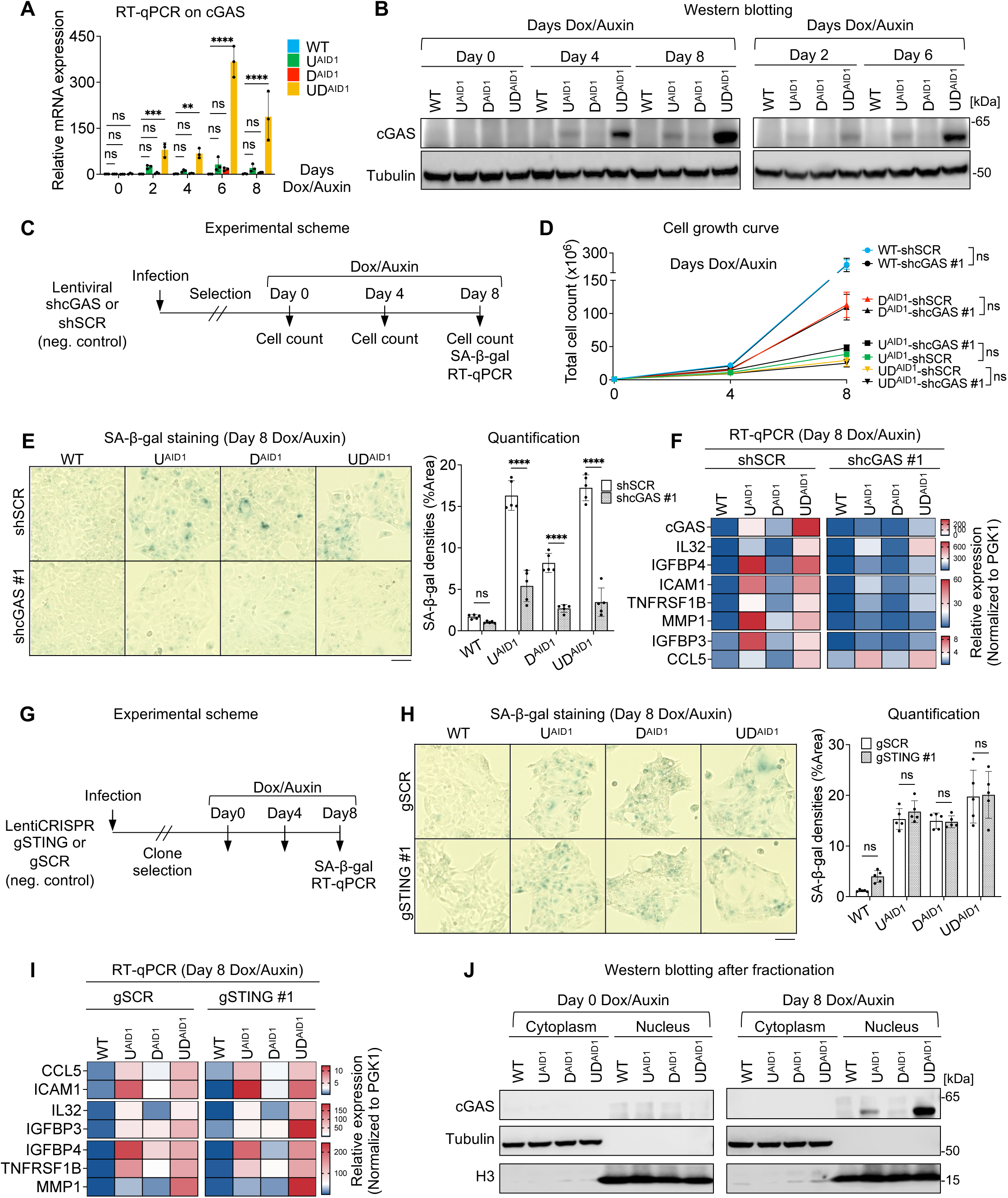
cGAS is necessary for SA-β-gal positivity and SASP expression during senescence caused by loss of DNA methylation, and it acts independently of STING. (A) Time-course RT-qPCR of cGAS gene expression in HCT116 lines upon Dox/Auxin treatment. N = 3 biological replicates. (B) Western blotting of cGAS protein in HCT116 lines at the indicated days upon Dox/Auxin treatment. (C) Scheme of cGAS knockdown experiments. (D) Total cell numbers at the indicated days upon Dox/Auxin treatment. N = 3 technical replicates. (E) SA-β-gal staining (left panel) and quantification (right panel) of the indicated lines. N = 5 fields of view. (F) RT-qPCR results of cGAS and selected SASP genes in the indicated conditions. (G) Scheme of STING knockout experiments. (H) SA-β-gal staining (left panel) and quantification (right panel) in the indicated conditions. N = 5 fields of view. (I) RT-qPCR results of selected SASP genes in the indicated conditions. (J) Western blotting of cGAS in cytoplasmic and nuclear fractions of HCT116 lines at the indicated days upon Dox/Auxin treatment. All scale bars are 50 µm. Data of (A), (D), (E) and (H) are presented as mean ± SD. Data of (A) are analyzed by two-way ANOVA with Dunnett’s multiple comparisons test. Data of (D), (E) and (H) are analyzed by two-way ANOVA with Sidak’s multiple comparisons test. Heatmap data of (F) and (I) are presented as mean from 3 technical replicates. In all figures we use the following convention: ** p < 0.01, *** p < 0.001, **** p < 0.0001, ns: non-significant.

To ascertain the potential role of induced cGAS during senescence induced by loss of DNA methylation, we performed an shRNA knockdown (Figure 5C). A cell-growth curve revealed no significant effect of cGAS knockdown on slow cell proliferation (Figure 5D). However, the results of the SA-β-gal assay showed a noticeable decrease in staining in the conditions of cGAS knockdown compared to the control (Figure 5E), and this finding was confirmed with an independent shRNA against cGAS (Figure S5A). RT-qPCR analysis confirmed the continuous suppression of cGAS after knockdown, and also revealed a downregulation of the SASP genes that are typically induced after DNMT1/UHRF1 depletion, suggesting that cGAS activation contributes to the SASP gene induction (Figure 5F).

Our next set of experiments addressed the possible role of STING in senescence induced by loss of DNA methylation. Our starting point was to examine the level of STING protein expression in the different cell types we used in our study. Western blotting showed that STING is indeed expressed in our HCT116 cells (Figure S5B), in agreement with previous findings (53). To explore its function, we employed CRISPR/Cas9 to generate STING knockout (KO) clones in each of the degron lines, and then assessed the senescence phenotype in these KOs (Figure 5G). The KO was efficient: no STING protein was detectable in any of the KO clones (Figure S5C). However, this genetic inactivation of STING had no effect on the induction of SA-β-gal positive cells (Figures 5H, S5D), nor on the expression of SASP genes (Figure 5I).

A STING-independent contribution of cGAS to senescence has been reported, but the mechanisms remain elusive (54). To test if cGAS is nuclear in our system, we performed a cellular fractionation on the degron cells after exposure to Dox/Auxin. We found that cGAS protein was accumulated in the nuclear fraction (Figure 5J). These data agree with a very recent report also describing that cGAS is nuclear when expressed in HCT116 cells (55).

Collectively, our results reveal that chronic DNA methylation loss after UHRF1/DNMT1 depletion leads to the induction of cGAS, which itself contributes to the activation of SASP genes and increased lysosomal activity. This role of cGAS occurs in the nucleus and is independent of STING.

### Prolonged DNA demethylation also triggers cancer cell senescence *in vivo*

Having established that the depletion of UHRF1 and/or DNMT1 triggers senescence in cancer cells *in vitro*, we next sought to test if this also occurred in an *in vivo* setting. To perform these experiments, we turned to the AID2 degron system, in which the OsTIR1 (F74G) derivative can be activated by micromolar concentrations of the auxin analog 5-Ph-IAA (33), with minimal side effects *in vivo* (33). We generated degron alleles of UHRF1 and/or DNMT1 in the HCT116 AID2 system, yielding the lines U^AID2^, D^AID2^ and UD^AID2^ (Figure 6A). We validated that 5-Ph-IAA treatment led to degradation of the tagged protein in these lines (Figure S6A), and consequently to a decrease of global DNA methylation (Figure S6B).

**Figure 6.**
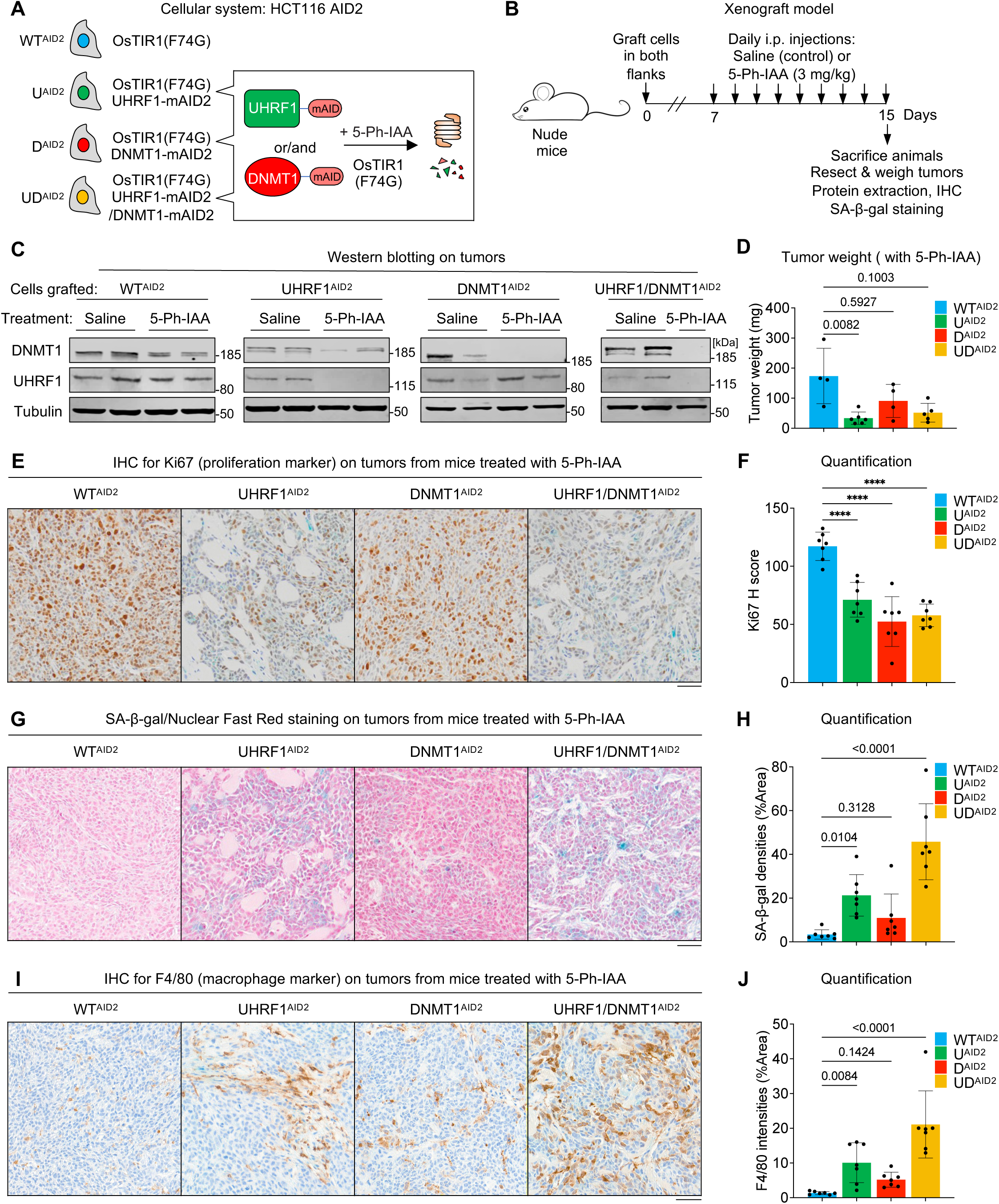
Senescence caused by loss of DNA methylation also occurs *in vivo*. (A) The auxin degron system in HCT116 cells constitutively expressing OsTIR1(F74G). (B) Scheme of the xenograft experiments. (C) Western blot validating the degradation of UHRF1 and/or DNMT1 upon 5-Ph-IAA treatment of mice. (D) Tumor weight assessment of the mice treated with 5-Ph-IAA. N = 4 tumors for WT^AID2^, 6 tumors for U^ADI2^, 4 tumors for D^AID2^, and 5 tumors for UD^AID2^. (E) Immunohistochemistry (IHC) staining of Ki67 in indicated conditions. (F) Quantification of Ki67 IHC using H score. N = 7 fields of view at 400× original magnification. (G) Representative images of SA-β-gal staining in indicated conditions. Nuclear Fast Red (NFR) was used for counterstaining. (H) Quantification of SA-β-gal staining. N = 7 fields of view at 300× original magnification. (I) IHC staining of F4/80 in indicated conditions. N = 7 fields of view at 400× original magnification. (J) Quantification of F4/80 IHC. All scale bars are 50 µm. All data are presented as mean ± SD. Data of (D), (H) and (J) are analyzed by Kruskal-Wallis test and Dunn’s multiple comparisons test. Data of (F) are analyzed by one-way ANOVA and Dunnett’s multiple comparisons test. We use the following convention: **** p < 0.0001.

Cells from each condition (HCT116 WT^AID2^, U^AID2^, D^AID2^ and UD^AID2^) were transplanted into the flanks of nude mice, and formed observable tumors after 7 days. We then administered either saline or 3 mg/kg of 5-Ph-IAA via intraperitoneal (i.p.) injections daily for an additional 8 days. At the end of the experiment the mice were sacrificed, and tumors were resected for a series of biological assessments (Figure 6B).

We first verified the *in vivo* efficiency of the AID2 system. In all tumors we examined, treatment with 5-Ph-IAA led to a complete degradation of the cognate target (Figure 6C), and this result was further confirmed by performing immunohistochemistry (IHC) for UHRF1 on tumor sections (Figures S6C, S6D). From these experiments we conclude that the AID2 system permits efficient depletion of UHRF1 and/or DNMT1 in our xenograft experiments.

We next ascertained the consequences of UHRF1 and/or DNMT1 depletion in the xenografts. The first parameter we measured was tumor mass. All four cell lines generated tumors of similar mass when mice were injected saline (Figure S6E); in contrast the treatment with 5-Ph-IAA, leading to protein degradation, caused the UD^AID2^ xenografts to be significantly smaller than control xenografts, with a similar trend also observed in D^AID2^ and U^AID2^, albeit failing to reach statistical significance (Figure 6D).

Our *in vitro* results led to the expectation that U^AID2^, D^AID2^ and UD^AID2^ cells, treated with 5-Ph-IAA *in vivo*, may proliferate less than control cells, and may enter senescence. We tested both predictions experimentally on tumor sections: the proliferation activity was estimated by Ki67 (a marker of proliferation) IHC, and the senescence by SA-β-gal staining.

Consistent with our *in vitro* findings, upon treatment, UHRF1 and/or DNMT1 degron xenografts showed a decreased signal of Ki67 compared to WT xenografts (Figures 6E, 6F), indicating a cell proliferation defect. We next performed whole-mount SA-β-gal assay on the tumors. After fixing and staining, tumor sections were counterstained with Nuclear Fast Red. Strikingly, the results revealed a dramatic increase of SA-β-gal positivity in U^AID2^ and UD^AID2^ xenografts, with a moderate increase of SA-β-gal positivity in D^AID2^ xenograft upon treatment (Figures 6G, 6H). In contrast, no SA-β-gal staining was detectable in xenografts from saline-treated mice (Figure S6F). Therefore, we conclude that loss of DNA methylation caused by degradation of UHRF1 and/or DNMT1, causes cancer cells to senesce *in vivo*.

The nude mice lack B and T lymphocytes, but they still possess components of the innate immune response, which can respond to a SASP (56). To investigate this in our *in vivo* model, we used IHC staining for F4/80, a marker of macrophages. The results showed an increase of F4/80 positivity in tumors depleted of UHRF1 or UHRF1 and DNMT1, supporting the notion that the resulting senescent cells promote inflammation (Figures 6I, 6J). Together with low proliferation and increased senescence, this macrophage response may be one of the factors contributing to the reduced growth of tumor cells lacking UHRF1 and/or DNMT1.

## Discussion

### Loss of DNA methylation induces senescence in multiple cancer cells

Through this study, we explored the consequences of lowering DNA methylation in cancer cells without promoting DNA damage. For this we exploited degron cell lines for UHRF1 and DNMT1, two enzymes required for DNA methylation maintenance. We found that the loss of DNA methylation without DNA damage induced senescence in cancer cells.

HCT116 cells deprived of UHRF1 and/or DNMT1 present 4 hallmarks of cellular senescence: withdrawal from the cell cycle, enlarged nuclei, expression of SASP, and increased SA-β-gal activity (Figures 1 and 2). Several arguments suggest that this phenotype is due to decreased DNA methylation, as opposed to other effects of removing UHRF1 and/or DNMT1. First, our rescue experiments with 9 point-mutants of UHRF1 and DNMT1 show a complete overlap between the ability to sustain DNA methylation and the ability to prevent senescence (Figure S1). Second, removing UHRF1 causes a more rapid loss of DNA methylation than removing DNMT1 (29), and it also causes a more rapid entry into senescence. Third, treatment with a last-generation inhibitor of DNMT1 that does not induce DNA damage (34) also causes cancer cells to become senescent (Figure 1). In parallel to our work, a degron approach has been recently applied by other authors to DNMT1 in DLD-1 colorectal cancer cells (57). This study shows that DNMT1 depletion leads to impaired proliferation with a G1-phase increase, S-phase decrease, and few sub-G1 cells; all of these observations are compatible with an entry into replicative senescence.

The senescence phenotype in cells with decreased DNA methylation is not restricted to a specific cell line, or a specific tumor type, as we have observed it in multiple cancer lines originating from the colon, lung, breast, and cervix (Figure 1). It will be instructive in future studies to determine if certain cancer lines deviate from this response and if so, why.

The senescence phenotype triggered by loss of DNA methylation is highly penetrant in HCT116 and other cancer cell lines, yet not reported in earlier publications. This likely results from prior studies using agents such as 5-aza-C or 5-aza-dC, which induce DNA damage, to research DNA demethylation. High doses of the drugs generate unsustainable DNA damage, resulting in apoptosis. Using smaller amounts of the drug may lead to a partial or delayed senescence phenotype, which could be challenging to detect if it is masked by non-senescent cells of if the observation period is not sufficiently long. Other studies have examined loss of DNA methylation after RNAi or shRNA approaches on UHRF1 or DNMT1 (58, 59, 35) and have not reported senescence. In retrospect, our results suggest that these approaches have likely imposed a positive selection on the cells with the least depletion, possibly obscuring the senescent phenotype. Finally, HCT116 cells that are genetically hypomorphic for DNMT1 (60, 61) are not reported to be senescent, but this could be because they retain a sufficient level of DNMT1 activity and/or because they have adapted genetically and/or epigenetically. In summary, we believe the degron approach has unmasked a phenotype that was difficult to detect with earlier tools.

Compared to previous strategies, the degron approach also permits more precise kinetic studies, allowing us to order in time the different elements of the cellular response to DNA demethylation. For instance, we report that, while the activation of an interferon response signature is an early event (within 2 days), the full induction of interferon lambda is more delayed (Figure 2). These observations pave the way for further studies in the future, to fill in additional mechanistic details underlying our findings.

The consequences of removing UHRF1 from normal cells differ from what we observe in cancer cells. For instance, genetically removing UHRF1 from liver hepatocytes stops their proliferation (62). However this phenotype is due to replication stress, DNA damage, and S-phase arrest (63, 64), all of which are absent in the degron cancer cells. In mouse neural stem cells, the genetic inactivation of UHRF1 does not affect proliferation, survival, or differentiation. Instead, it induces the reactivation of intracisternal type A particle (IAP) repeats, eventually leading to apoptosis (65). To extend our work in the coming years, a broad endeavor will be to explore what mechanisms underlie the different responses of normal and cancer cells to the loss of DNA methylation, and to determine if our findings could relate to the changes of DNA methylation that occur in normal cells during organismal aging (66).

### Non-canonical mechanisms underpin senescence induced by loss of DNA methylation

Senescence induced by loss of DNA methylation differs in several ways from other instances of senescence previously reported in normal and cancer cells.

First, many of the known senescence-inducing stresses, such as oncogenic activity, DNA damage, or metabolic stresses, converge on p53 and/or Rb (67), but these two factors are dispensable for loss of DNA methylation to trigger senescence (Figure 3).

Another salient result is that DNA damage, which plays a central role in many senescence types, is unlikely to be involved in senescence triggered by DNA demethylation (Figure 3). Given that high levels of retrotransposition cause DNA damage (68), this result also implies that retrotransposition is either mild or absent in cells with decreased DNA methylation. This may be explained in part by the observation that DNA damage itself can activate repeated elements (69). Therefore, a 5-azacytidine treatment would cause a stronger reactivation of repeated elements than DNA demethylation alone.

An additional difference is that, in cells with lower DNA methylation, cytosolic p21 plays a protective role against apoptosis (Figure 4), at variance with nuclear p21 inhibiting the cell cycle in canonical senescence models (67).

A last distinction concerns the role of cGAS. Cytosolic chromatin fragments can trigger senescence (50, 51), a phenomenon implying cytosolic cGAS, which becomes catalytically active and produces cyclic GMP-AMP (cGAMP), thereby stimulating STING. In cells with decreased DNA methylation, cGAS is nuclear and STING is not required for senescence (Figure 5), so different processes must be at play. In particular, the results argue that cytoplasmic DNA originating from transposable elements do not have a major contribution to the phenotypes we observe, unlike what is seen in replicative senescence (68). A recent study found that chromatin-bound cGAS can recruit the SWI/SNF complex at specific chromatin regions to regulate gene expression (55), thus a similar process might be at play in our system. It has also been reported that UHRF1 inhibits the activity of nuclear cGAS (70), and this may also contribute to our observations.

New chromatin-based mechanisms underlying SASP expression and senescence are regularly being discovered. For instance, spurious intragenic transcription participates to senescence (71), while the histone chaperon HIRA plays a histone-independent role in SASP induction (72). In follow-up studies, it will to determine if these phenomena contribute to the senescence phenotypes we observed upon DNA methylation decrease.

### Therapeutic implications

Our xenograft experiments (Figure 6) show that cancer cells with decreased DNA methylation undergo senescence *in vivo*. Inducing senescence in tumor cells is emerging as a promising therapeutic avenue (67, 73). Not only does this strategy suppress their proliferation, but the accompanying SASP also has a paracrine pro-senescent effect on neighboring tumor cells that may have escaped treatment. Two added benefits are that the SASP can make the vasculature more permeable to drugs (74), and that it activates an inflammatory response that can promote clearing by the immune system (73). Nevertheless, inducing senescence in tumors may be a double-edged sword, as senescent cells can become immunosuppressive, induce an Epithelium to Mesenchyme Transition (EMT) in non-senescent cells, or promote angiogenesis (73). It may be possible to circumvent these problems by combining the induction of senescence by loss of DNA methylation with a senolytic treatment, to effectively clear the senescent cancer cells (73).

Several existing cancer therapies can be used to promote cancer cell senescence, including the use of genotoxics (75). Our data show that an additional option to achieve this goal is to decrease DNA methylation, without causing DNA damage. A clinically important implication of our results is that this strategy should be applicable to the many tumors lacking p53 and/or Rb.

Molecularly, decreasing DNA methylation without DNA damage can be achieved with DNMT1 inhibitors that do not incorporate into DNA, such as the recently described GSK3685032 (34), or its derivatives (76). However, our degron experiments show that UHRF1 inhibition has a faster and/or stronger effect than DNMT1 inhibition, which likely stems from DNMT1-independent roles of UHRF1 (29). This suggests that the development of UHRF1 inhibitors may be a promising strategy for inducing senescence without DNA damage in cancer cells. Our work identifies domains of UHRF1 that are essential for preventing senescence of cancer cells (Figure S1). These domains are potentially druggable, and UHRF1 has already been used for chemical screens *in vitro* (77–80), but an efficient and specific inhibitor has yet to emerge. In some instances, targeted protein degradation by a proteolysis targeting chimera (PROTAC) is superior to chemical inhibition (81), and this approach could be explored for UHRF1 as well.

As for any other drug, a prerequisite for UHRF1 inhibitors to be useful is the possibility to treat cancer cells without harming healthy cells. This might be solved by addressing the drugs specifically to the tumor. Alternatively, given that many tumors highly overexpress UHRF1 (35, 82), they might possibly have developed a higher reliance on UHRF1 than normal cells, which would open a possible therapeutic window. These and other questions can be addressed in the future on the basis of the findings reported here.

## Materials and Methods

### Cell lines and cell culture

The HCT116 cells are male. Their derivative expressing OsTIR1 under a doxycyclin-inducible promoter was developed and published by the Kanemaki lab (30), and we described in a previous publication how we used them to generate UHRF1, DNMT1, and UHRF1/DNMT1 degron cells, and how we rescued these cells genetically with WT or mutant constructs (29). We also used DLD1 cells (male) constitutively expressing the OsTIR1(F74G) improved variant (33) to generate degron alleles of UHRF1, DNMT1, or both, in this AID2 system (29).

For the current publication, we generated a new series of degron lines, in the HCT116 AID2 background. The cells were transfected with CRISPR-Cas9 and donor plasmids using Lipofectamine 2000. After 700 µg/mL G418 or 100 µg/mL hygromycin selection, we obtained HCT116 AID2 lines (UHRF1-mAID2, DNMT1-mAID2 and UHRF1-mAID2/DNMT1-mAID2). Other human cancer cell lines including HT29 (female), A549 (male), MCF7 (female) and HeLa (female) were acquired from the ATCC.

To achieve degradation of the AID-tagged protein, AID1 cells were seeded and incubated with 2 µg/mL doxycycline (Dox) and 20 µM auxinole for 24 hr. Afterwards, the medium was replaced with a fresh medium containing 2 µg/mL Dox and 500 µM indole-3-acetic acid (IAA, auxin). For AID2 cells, they were seeded one day prior to the addition of medium containing 1 µM 5-Ph-IAA, an IAA derivative. The medium with either Dox/Auxin or 5-Ph-IAA was replaced every two days.

HCT116 and HT29 cells were cultured in McCoy’s 5A medium supplemented with 10% FBS, 2 mM L-glutamine, 100 U/mL penicillin and 100 μg/mL streptomycin. DLD1 cells were cultured in RPMI-1640 medium with the same supplements as above. A549, MCF7 and HeLa cells were cultured in DMEM medium also supplemented identically. All cultures were incubated at 37 °C under a humidified atmosphere containing 5% CO_2_. Cells were tested monthly for mycoplasma infection. The cell identity was confirmed by short tandem repeat (STR) analysis.

As a positive control for apoptosis induction, HCT116 WT cells were treated with 40 µg/mL 5-Fluoro-Uracil (5-FU) for 2 days. As a positive control of DNA damage induction, WT were treated with 2 mM hydroxyurea (HU) or 10 µM etoposide (Eto) for 4 hr.

### Plasmid construction and viral infection

shRNA targeting Scramble (negative control), cGAS or p21 were cloned into the pLKO.1-blast vector (Addgene #26655). Plasmids were generated using PCR, restriction enzymes, or Gibson Assembly (NEB) Cloning techniques. The oligonucleotide sequences inserted into the pLKO.1-blast vector are available in Supplementary Table 1. The vector pLenti X2 Blast/shp16 (w112-1) (Plasmid #22261) and pLenti X2 Blast/pTER shLUC (w618-1) (Plasmid #20962) were from Addgene.

Human papillomavirus type 16 (HPV16) E6-E7 combined genes were amplified from the template vectors (pLXSN E6-E7, Addgene #52394) and then cloned into pLenti6.2/V5-DEST lentiviral vector (Invitrogen). gRNA targeting STING and control are gifts from Nicolas Manel (Institute Curie, FR). All plasmids underwent sequencing prior to their utilization.

For viral infections, HCT116 WT and degron lines were plated overnight, then infected with retroviruses in the presence of 6 µg/mL polybrene (Sigma-Aldrich) for 16 h. After 48 h, infected cells were selected with 10 µg/mL blasticidin for 7 days.

### Cell counting

Cells were initially seeded at 0.125 million per well in 6-well plates and cultured in medium supplemented with 2 µg/mL Dox and 20 µM auxinole. On the following day (Day 0), the medium was refreshed with one containing 2 µg/mL Dox and 500 µM auxin. Cell counts were performed on Day 4, after which cells were passaged at a density of 0.25 million, and again on Day 8. The Dox/Auxin-containing medium was renewed every two days. A BioRad TC20 automated cell counter was employed to determine total cell numbers.

### Protein extraction

Cells were harvested after trypsinization, washed twice with PBS, and lysed with RIPA buffer (Sigma-Aldrich) with protease inhibitor cocktail (Roche) and phosphatase inhibitor cocktail 3 (Sigma-Aldrich) for 20 min on ice, and then sonicated with series of 30 s ON / 30 s OFF for 5 min on a Bioruptor device (Diagenode) and centrifuged at 16,000 g for 15 min at 4 °C. The supernatant was collected and quantified by Pierce BCA protein assay kit (Thermo Fisher Scientific).

### Cell fractionation

Cells pellets (5 million) were gently lysed in 500 μL of hypotonic buffer (20 mM of Tris-HCl (pH 7.6), 3 mM of MgCl_2_, 10 mM of NaCl) supplemented with protease inhibitor cocktail (Roche) and phosphatase inhibitor cocktail 3 (Sigma-Aldrich). Incubate on ice for 20 min. Add 25 μL detergent (10% NP-40) and vortex for 10 seconds at highest setting. Following centrifugation for 10 min at 3,000 rpm at 4 °C, the supernatants were collected as the cytoplasmic fraction. Pellets were washed twice with hypotonic buffer and then were lysed by 200 μL RIPA buffer. After incubation on ice for 30 min, the lysates were centrifuged for 30 min at 13,000 rpm at 4 °C. The supernatants then were collected as the nuclear fraction.

### Western blot

Equivalent amounts of protein extract per sample were mixed with NuPage 4X LDS Sample Buffer and 10X Sample Reducing Agent (Thermo Fisher Scientific) and denatured at 95°C for 5 min. Samples were resolved on a pre-cast SDS-PAGE 4-12% gradient gel (Invitrogen) and transferred to a nitrocellulose membrane (Millipore). The membranes were blocked in 5% (w/v) non-fat milk in PBST buffer (PBS with 0.1% Tween 20) or 5% (w/v) bovine serum albumin (BSA) in TBST buffer (TBS with 0.1% Tween 20), then incubated overnight at 4°C with the appropriate primary antibodies diluted in 5% milk PBST or 5% BSA TBST. After washing with PBST or TBST, the membranes were treated with fluorophore-linked or HRP-conjugated secondary antibodies and visualized using the LI-COR Odyssey-Fc imaging system (LI-COR). All the antibodies are listed in Supplementary Table 1.

### BrdU labeling and flow cytometry

To analyze the cell cycle, HCT116 cells were seeded in a 6-well plate and treated with Dox/Auxin for up to 8 days. The cells collected at Day 0, Day 4 and Day 8 were treated with 10 μM 5-bromo-2’-deoxyuridine (BrdU) (Sigma-Aldrich) for 1 hour at 37 °C in a CO_2_ incubator. Cells were washed, trypsinized, and resuspended in 750 µL of PBS before adding 2250 µL of ice-cold ethanol for fixation (yielding a final volume of 3 mL with 75% ethanol). The fixed cells were incubated for at least 30 min at −20 °C prior to flow cytometry analysis. Samples were denaturated in 2 N HCl for 15 min at room temperature, followed by two washes in PBS + 1% BSA. Samples were then incubated in 200 µL of anti-BrdU-FITC antibody (BD Bioscences) in PBS + 1% BSA for 60 min. After a washing step, samples were then incubated in PBS containing propidium iodide (PI) (1:500, Invitrogen) with 150 µg/mL RNAse A (Qiagen) overnight at 4 °C in the dark. Cell cycle profiles were measured by flow cytometry using the Cytoflex SRT (Beckman Coulter). Data were analyzed with the FlowJo v10.4 software. Cell populations were gated by size using SSC-A and FSC-A parameters. For single-cell resolution, we employed FSC-H versus FSC-A gating. All gated information was supplied for cell population analysis.

### TUNEL assay

HCT116 cells were seeded on coverslips two days before fixation. The cells were fixed with 4 % paraformaldehyde for 15 min then permeabilized using 0.25 % Triton X-100 in PBS for 20 min at room temperature. For the TUNEL assay, the following steps were performed using the Click-it Plus TUNEL Assay according to the manufacturer’s protocol (Invitrogen). For a positive control, cells were treated with DNase I (Invitrogen) for 30 min at room temperature. Cells were additionally stained for DNA using 1× Hoechst 33342 (Life Technologies). Cells were imaged using Leica DMI6000 (Leica Microsystems).

### Annexin V assay

Cells adhered to the dishes and those detached in the supernatant were collected for apoptosis assays. Cell pellets were then resuspended in Annexin Binding buffer (10 mM HEPES, 140 mM NaCl, 2.5 mM CaCl_2_ pH 4). The following steps were performed using the Annexin V Conjugates for Apoptosis Detection kit (Invitrogen). Cells were stained with 0.1% (v/v) Annexin V and 1× Hoechst 33342 (Life Technologies) solution. Stained cell suspensions were incubated in the dark for 15 min at room temperature. 400 µL of 1× binding buffer was added to each tube and samples were then immediately analyzed by flow cytometry using the Beckman Coulter Cytoflex SRT.

### Immunofluorescence

Cells were fixed in 4% paraformaldehyde at room temperature for 15 min. After fixation, cells were permeabilized with 0.5% Triton X-100 in PBS for 10 min at 4 ℃, then washed with PBS. Cells were blocked with 1% BSA in PBS at room temperature for 30 min, then incubated with γ-H2AX antibody (Sigma-Aldrich 05-636, 1:1,000) for 1 hr at room temperature. After washing 3 times with PBST, cells were incubated with secondary antibodies conjugated with Cy5 for 1 hr at room temperature, washed with PBST 3 times, and finally mounted with ProLong Diamond Antifade Mountant with DAPI (P36961, Thermo Fisher Scientific). Images were obtained using Leica DMI6000 (Leica Microsystems) and MetaMorph Leica v6.1 software.

### RNA extraction and quantitative reverse transcription PCR (RT-qPCR)

Total RNA was extracted from cells with the RNeasy Plus Mini kit (Qiagen) according to the manufacturer’s instructions and quantified using Nanodrop or Qubit RNA BR Assay kit on the Qubit 2.0 Fluorometer (Thermo Fisher Scientific). 1 µg of total RNA was reverse transcribed using SuperScript IV Reverse Transcriptase (Thermo Fisher Scientific) and random primers (Promega). RT-qPCR was performed using Power SYBR Green (Life Technologies) on Applied Biosystems QuantStudio 6 Pro. Housekeeping gene PGK1 was used for normalization. All the primer sequences for RT-qPCR are listed in the Supplementary Table 1.

### Senescence-associated β-galactosidase (SA-β-gal) staining

SA-β-gal staining was performed with the Senescence Cells Histochemical Staining Kit (Sigma– Aldrich, cat # CS0030-1KT). Briefly, monolayered cells in 6-well plates were incubated with fixation buffer containing 2 % formaldehyde for 7 minutes at room temperature, followed by 37 °C overnight incubation in staining solution, supplemented with 1 mg/mL of 5-bromo-4-chloro-3-indolyl-β-d-galactopyranoside (X-gal). Plates with staining solution were kept overnight at 37 °C without CO_2_ until the cells were stained blue. The plates were sealed with parafilm to prevent them from drying out. The stained cell images were taken using LEICA DMi1 (Leica Microsystems).

### Conditioned medium (CM) collection, CM culture and enzyme-linked immunosorbent assay (ELISA)

Conditioned media from four conditions were prepared. Conditioned medium from HCT116 AID1 cells: WT and degron cells were cultured in McCoy’s 5A medium supplemented with 10% FBS and Dox/auxin for 8 days. Cells were then washed twice with PBS, followed by incubation in FBS-free McCoy’s 5A medium for 20 hr. Conditioned medium from the interferon response positive control: HCT116 WT cells transfected with PolyI:C (1 µg/mL, Invivogen) using Lipofectamine RNAiMAX (Invitrogen) for one day underwent a wash step and medium replacement before incubating in FBS-free McCoy’s 5A medium for another 20 hr. Conditioned medium from interferon response negative control: WT cells were cultured in McCoy’s 5A medium without FBS for 20 hr. CM from DNA damage induced senescence: WT cells were treated with 1 µM etoposide for 24 hr, followed by a 3-day fresh medium culture before switching to FBS-free McCoy’s 5A medium for an additional 20 hr.

Serum-free conditioned media was collected after counting cell numbers to ensure equal volumes from equivalent cell counts. Media was centrifuged at 300 x g for 5 min to remove cellular debris and filtered through a sterile 0.2 µm filter (Thermo Fisher Scientific).

For CM culture, WT cells were culture in CM supplemented with an additional FBS for 24 hr. For ELISA, CM from WT, PolyI:C transfected WT, Dox/Auxin treated WT and degron cells was 10× concentrated using Amicon Ultra centrifugal filter (Millipore). The ELISA assay was performed according to the manufacturer’s protocol (Thermofisher, cat #88-7296) to detect the presence of human IL-29 (Interferon lambda 1). Detection was performed using AMR-100 Microplate Reader at 450 nm (Hangzhou Allsheng).

### Genomic DNA extraction and Luminometric methylation (LUMA) assay

Genomic DNA was extracted by Monarch Spin gDNA extraction kit following the manufacturer’s protocol. LUMA analysis was conducted following standard procedures. In brief, to assess global CpG methylation, 500 ng of genomic DNA was digested with Msp I + EcoR I and Hpa II + EcoR I (New England Biolabs) in parallel reactions. EcoR I was included as an internal reference. CpG methylation percentage is defined as the Hpa II/Msp I ratio. Samples were analyzed using PyroMark Q24 Advanced pyrosequencer (Qiagen).

### Xenograft Mouse models

Six week old Rj:NMRI-Foxn1 nu/nu female mice were acquired from Janvier Labs and maintained under standard conditions (standard diet and water ad libitum) at 23 °C with 12 h light and 12 h dark cycles, in a specific pathogen-free (SPF) animal facility; they were acclimated a week before performing experiments. To form xenografts, a suspension of HCT116 cells in 50% Matrigel (Corning #354234) was injected subcutaneously into the right and left flanks of each mouse. Large (L) and small (l) length of each tumor were measured using a caliper and the tumor volume was calculated using the following formula: Volume = ½ (L × l^2^). Before drug treatment, we first let the tumors engraft until they reached a volume of approximately 60 mm^3^. The mice were injected intraperitoneally daily for 8 days with 5-Ph-IAA (3 mg/kg) or with NaCl 0.9 %. Animal work was conducted according to the ethical guidelines applicable in France, in a duly authorized animal facility (Plateforme du Petit Animal du CRCL (P-PAC), agreement number D693880202). Our project received the agreement number APAFIS #49702-202404181412449 v3 from the French Ministry of Education and Research.

### Tumor protein extraction

Tumors collected from the xenograft mouse models were cut and placed in a RIPA buffer (Sigma-Aldrich) with protease inhibitor cocktail (Roche) and 0.05% PMSF. Samples were further homogenized using Precelly’s ceramic beads 1.4 mm (13113–325) and following the manufacturer’s protocol for protein extraction with the Precelley’s 24 Dual Lysis and Homogenization (P002511-PEVT0-A.0, Bertin Technologies).

### Whole-mount SA-β-gal staining

At resection, tumors were snap frozen and kept at −80°C until the assay. After thawing, tumors were fixed and SA-β-gal whole mount staining was performed overnight according to the procedures described in the Senescence Beta-Galactosidase Cell Staining Kit (Cell Signaling Technology #9860). All incubations and washes were performed with fresh and filtered solutions. Stained tissues were then washed twice in PBS and post-fixed with 4% paraformaldehyde before paraffin embedding for sectioning. 8 µm sections were cut and counterstained with Nuclear Fast Red (Sigma-Aldrich #N30-20-100mL).

### Immunohistochemistry (IHC) staining and scoring

For IHC staining, 4 µm fixed tumor sections were incubated for 15 min at 95 °C in a solution of deparaffinization and antigen retrieval at pH 6 or pH 9. Then they were rinsed and incubated 20 min at room temperature with primary antibodies against UHRF1, Ki67 or F4/80. Details of the antibodies used are provided in the Supplementary Table 1. The revelation in brightfield was performed by “Bond Polymer Detection”, (Leica DS9800). Sections were lightly counterstained with hematoxylin, then dehydrated and cleared in graded alcohol and Ottix plus (MM-France), and finally covered with glass slips.

An IHC score was calculated by summing the products of staining intensities (0–3) and percentage of stained cells in each intensity (0–100); H scores ranged from 0 to 300. The intensity level was evaluated by the IHC profiler plugin in ImageJ software (v2.14.0). 0 point was assigned for negative, 1 for low positive, 2 for positive and 3 for high positive. Each section was assayed for seven independent high magnification (×400) fields to get the average scores.

### RNA sequencing, data processing and Virtual Inference of Protein-activity by Enriched Regulon analysis (VIPER) (46)

Sequencing libraries were made from poly-A RNA, as recommended by Illumina, and sequenced using either an Illumina NovaSeq 6000 sequencer in Novogene Co., Ltd. RNA-seq 150bp paired-end reads were assessed for quality using the ‘FastQC’ algorithm, trimmed as appropriate using the algorithm ‘trim-galore’ (version 3.0), then the reads were aligned to the human hg38 reference genome using STAR (v2.6.1d). FeatureCounts (v1.5.0-p3) was used to count the read numbers in each gene before differential expression analysis using the linear modeling tool DESeq2 was performed. Using both differential expression methods, significantly changing expression was defined as an adjusted p value (FDR) < 0.05. The expression levels were normalized by transcript per million (TPM) values. Gene ontology analysis was performed using Gene Set enrichment Analysis (GSEA) software (v4.1.0). msVIPER function in viper R package was used for protein activity analysis. The human colon adenocarcinoma context-specific regulatory network was obtained from aracne.networks R package. This package contains ARACNe-inferred networks from TCGA tumor datasets.

### Statistical Analysis and visualization

Results were statistically analyzed in GraphPad Prism (v10.3.1, GraphPad Software Inc., San Diego, CA) and presented as mean ± SD or mean. Statistically significant differences between two groups were assessed by two-tailed unpaired t-test. Comparisons between two groups were assessed using the unpaired Student’s t-test (2-tailed). Comparisons between more than two groups were assessed by one-way ANOVA or Kruskal-Wallis test, followed by multiple comparisons test. p < 0.05 was considered significant. ns, not significant; *p < 0.05; **p < 0.01; ***p < 0.001; ****p < 0.0001 for indicated comparisons. Statistical details of each experiment can be found in the Results and Figure Legend sections. Principal component analysis (PCA), differentially expressed gene analysis, hierarchical clustering analysis and heatmap were generated by R software (v4.1.0).

## Acknowledgments

This work was supported by the following grants: Agence Nationale de la Recherche (ANR-23-CE12-0015-01, to PAD and DB) and Institut National du Cancer (INCA_18350, to PAD). The team of NM is supported by Fondation pour la Recherche Médicale (EQU202103012774), Agence Nationale pour la Recherche (ANR-21-CE15-0037-01 and ANR-21-CE13-0030-02), Fondation ARC (ILUMINAGE), Sidaction (22-2-AEQ-13411), and the Clayco Foundation. XC received support from “Fondation pour la Recherche Médicale” for the last year of her PhD (FDT202404018544). KY was supported by ARC foundation CDD Post-doctorant en France (Ref. No. PDF20181208337), Labex “Who Am I?” postdoctoral support, and a JSPS Postdoctoral Fellowship for Research Abroad (Ref. No. 202260432). The work of FM and TI was supported by Research Support Project for Life Science and Drug Discovery (Basis for Supporting Innovative Drug Discovery and Life Science Research (BINDS)) from AMED under Grant Number JP23ama121022. The authors are grateful to the ImagoSeine core facility of Institut Jacques Monod, member of France-BioImaging (ANR-10-INBS-04), with the support of Plan Cancer, Region Ile-de-France and Fondation Bettencourt Schueller (R03/75-79). We also thank the Vectorology platform, Epigenetics platform, Microscopy platform and Bioinformatics/Biostatistics Core Facility (BIBS) at the CNRS Epigenetics and Cell Fate Unit (Université Paris Cité), for providing access and technical advice. We are especially grateful to the Hist’IM facility (Institut Cochin) for excellent histology support.

We are very indebted to Saadi Khochbin, Allison Bardin, Gael Cristofari, Corinne Abadie, Bill Keyes, Julien Cherfils-Vicini, Vjekoslav Dulic, Clément Hua, Nadine Laguette, Han Li, Raphael Margueron, François Radvanyi, Laurent Reber, and David Roulois for useful advice. We thank the following colleagues for the gift of useful reagents: Fadila Rayah, Catherine Postic, Emma Lamana, and Pierre Bruhns.

## Author contributions

XC, KY and PAD conceived, designed and supervised the project.

XC, KY, and BR established the constructs and cell lines used in this study.

XC, KY, and BR performed molecular and cellular biology experiments, collected and analyzed data, with the assistance of MN and LF.

XC analyzed, interpreted and visualized the RNA sequencing data.

DG and DV performed xenograft experiments, with the assistance of N. Martin.

XL assisted with experiments related to cGAS and STING.

XC and PD performed histology analyses.

FM performed experiments related to DNA methylation.

XC and BR wrote the original draft of the manuscript. PAD reviewed and edited the draft.

TI, N. Manel, MTK, DB and PAD acquired funding.

---

**Figure S1.**
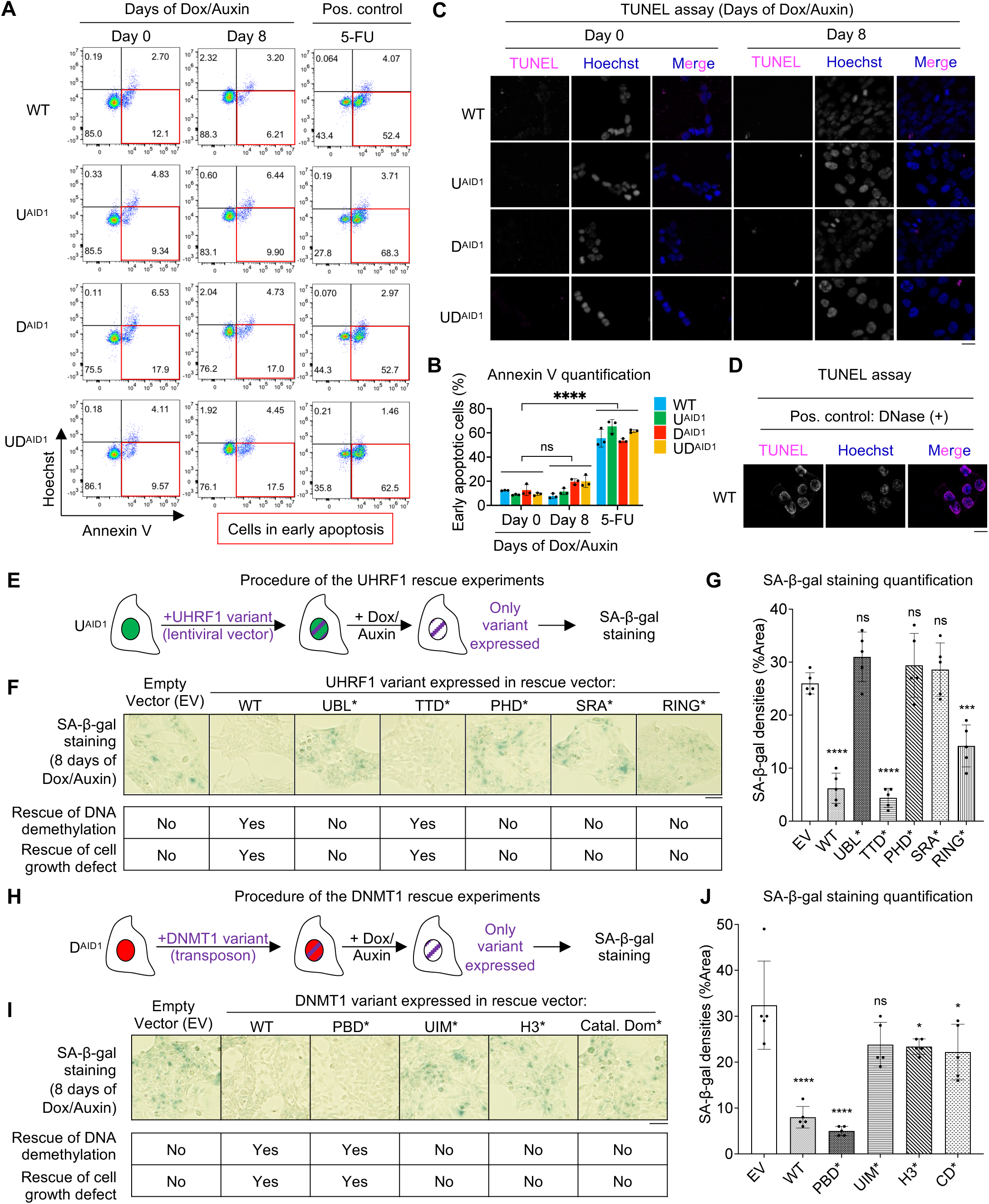
Lack of apoptosis in cells depleted of UHRF1 or DNMT1; rescue experiments support the role of DNA methylation maintenance in preventing senescence. (A) Representative results of Annexin V/Hoechst 33342 double staining upon Dox/Auxin treatment or 5-Fluoro-Uracil (5-FU, 40 µg/mL, 2 days) treatment as positive control. Cells in early apoptosis are indicated by the red square. (B) Percentages of early apoptotic cells (Annexin V positive/Hoechst 33342 negative). N = 3 biological replicates. (C) Representative TUNEL assay images upon Dox/Auxin or 5-FU (40 µg/mL, 2 days) treatment. Scale bar: 20 µm. (D) Positive control of TUNEL assay. Scale bar: 20 µm. (E) Schematic of UHRF1 rescue experiment. U^AID1^ cells were stably rescued with wild-type UHRF1 (WT) or point mutants inactivating the indicated domains. (F) Representative images of SA-β-gal staining upon 8-day auxin treatment with endogenous UHRF1-depleted HCT116 cells complemented by the indicated mutants. Scale bar: 50 µm. (G) Quantification of SA-β-gal staining. N = 5 fields of view. (H) Schematic of DNMT1 rescue experiment. D^AID1^ cells were stably rescued with either wild-type DNMT1 or point mutants inactivating the indicated domains. (I) Representative images of SA-β-gal staining upon 8-day auxin treatment with endogenous DNMT1-depleted HCT116 cells complemented by the indicated mutants. Scale bar: 50 µm. (J) Quantification of SA-β-gal staining. N = 5 fields of view. All data are presented as mean ± SD. Data of (B) are analyzed by two-way ANOVA test with Dunnett’s multiple comparisons test. Data of (G) and (J) are analyzed by one-way ANOVA test with Dunnett’s multiple comparisons test. In all figures we use the following convention: * p < 0.05, *** p < 0.001, **** p < 0.0001, ns: non-significant.

**Figure S2.**
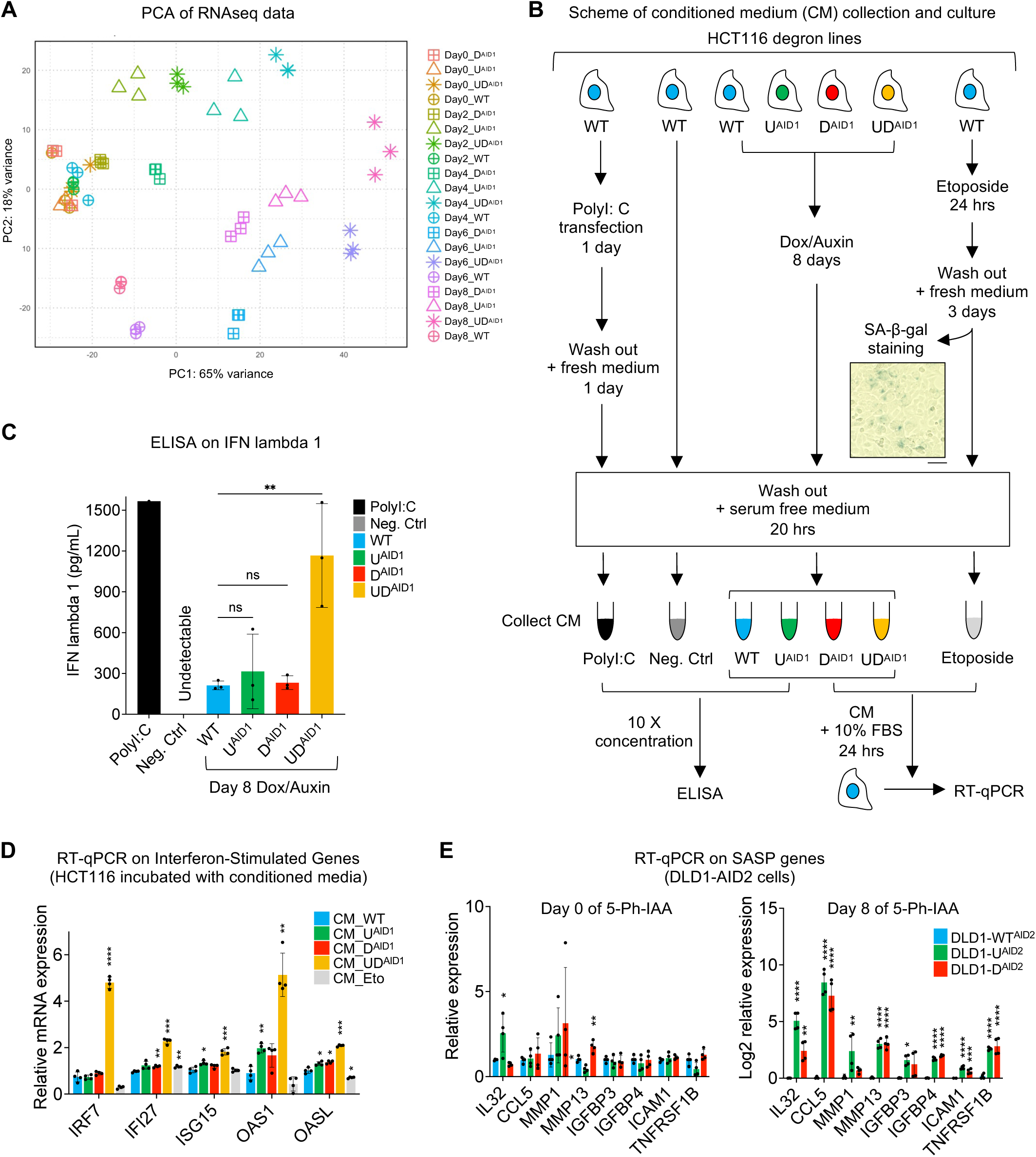
PCA analysis of the RNA-seq data, experiments with conditioned medium, RT-qPCR validations. (A) Principal component analysis (PCA) of RNA-seq data. B) Scheme of experiments with conditioned medium (CM) and validation of etoposide (1 µM) induced senescence by SA-β-gal staining. Scale bar is 50 µm. (C) Enzyme-linked immunosorbent assay (ELISA) on interferon (IFN) lambda 1 in the serum-free medium of HCT116 lines upon Dox/Auxin treatment at Day 8 or HCT116 WT transfected with PolyI:C (1 µg/mL) as positive control or WT without Dox/Auxin treatment as negative control. N = 3 biological replicates in WT, U^AID1^, D^AID1^ and UD^AID1^. (D) RT-qPCR of selected interferon stimulated genes (ISGs) expression in WT cultured with indicated conditioned medium for 1 day. N = 3 technical replicates. (E) Relative expression of selected SASP genes in DLD1 AID2 degron cells at Day 0 (left) and Day 8 (right) of 5-Ph-IAA treatment. N = 2 biological replicates and 2 technical replicates. Data of (B), (D) and (E) are presented as mean ± SD and analyzed by one-way ANOVA test with Dunnett’s multiple comparisons test. We use the following convention: * p < 0.05, ** p < 0.01, *** p < 0.001, **** p < 0.0001, ns: non-significant.

**Figure S3.**
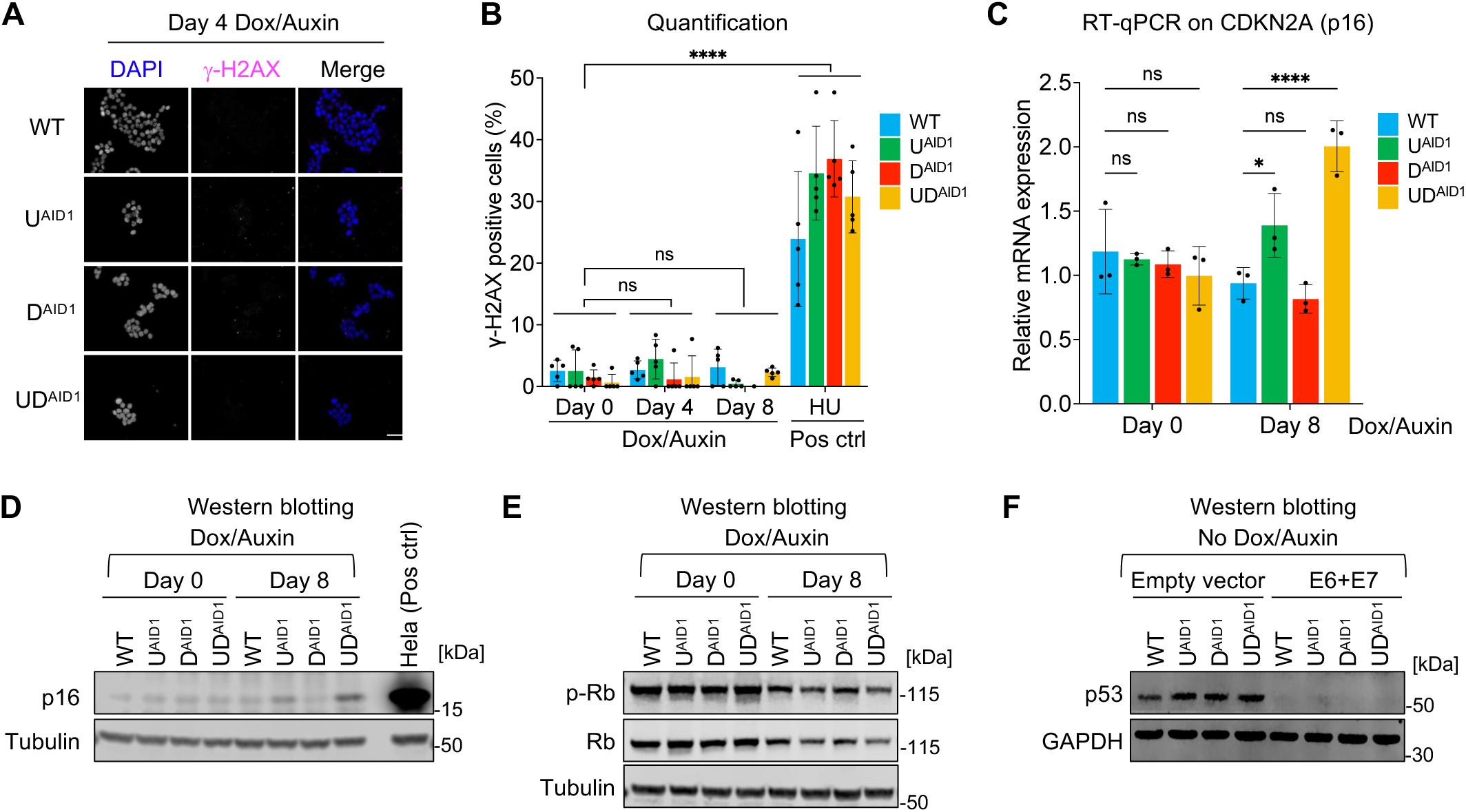
DNA demethylation-induced senescence is independent of p53 and p16/Rb: further controls and quantifications. (A) Immunofluorescent staining for γ-H2AX in indicated HCT116 lines upon Dox/Auxin treatment at Day4. Scale bar: 50 µm. (B) Quantification of the percentage of γ-H2AX foci positive cells. N = 5 fields of view. (C) qRT-PCR results of CDKN2A (p16) in indicated conditions. N = 3 biological replicates. (D) Immunoblots for p16 in indicated HCT116 lines upon Dox/Auxin treatment. Hela cells used as positive control. (E) Immunoblots for total Rb and phosphorylated Rb (Ser807/811) in indicated conditions. (F) Immunoblots for p53 in indicated HCT116 lines upon Dox/Auxin treatment. Data of (B) and (C) are presented as mean ± SD and analyzed by two-way ANOVA with Dunnett’s multiple comparisons test. In all figures we use the following convention: * p < 0.05, **** p < 0.0001, ns: non-significant.

**Figure S4.**
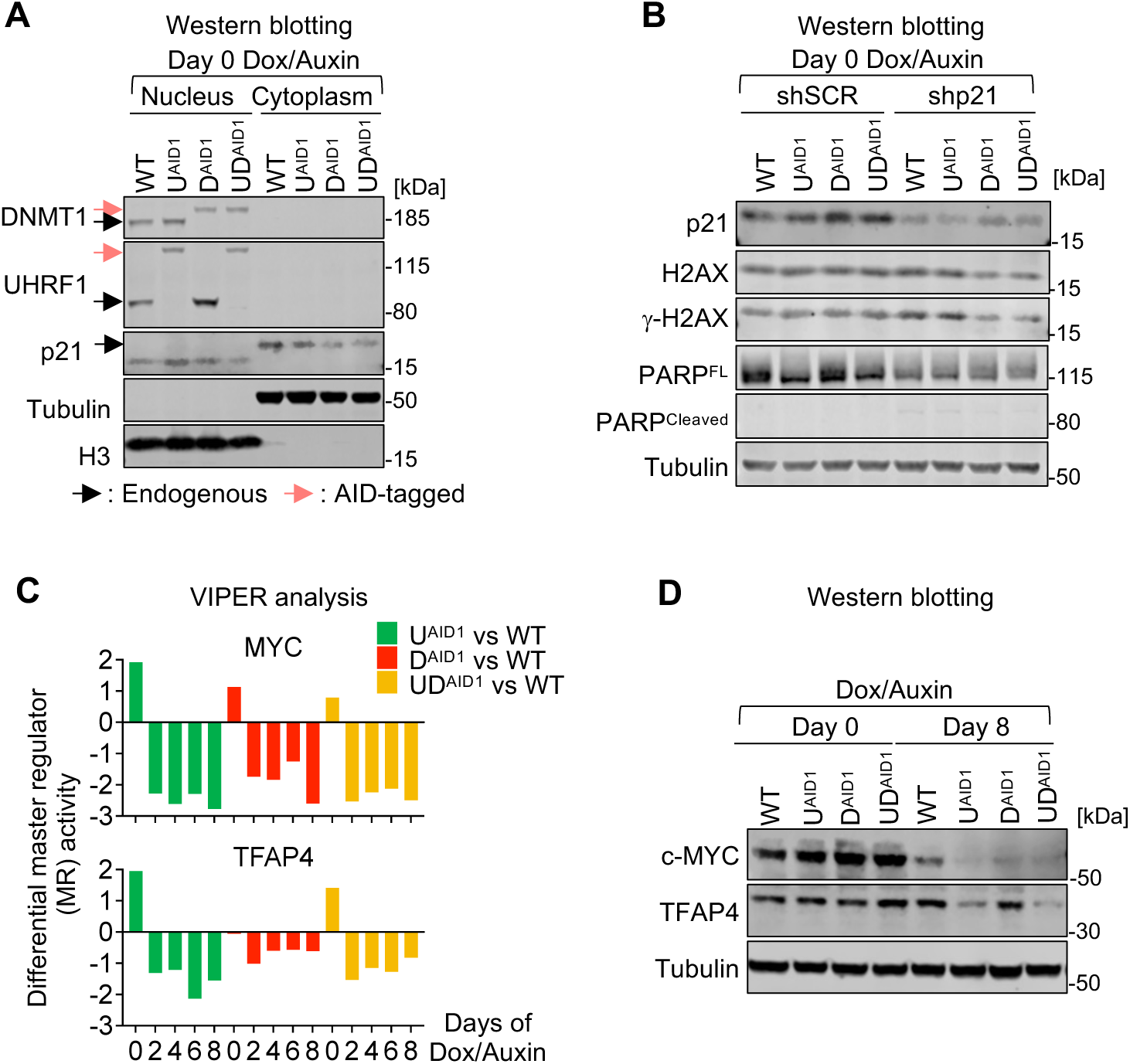
Further investigation of the MYC-TFAP4-p21 cascade. (A) Immunoblots for DNMT1, UHRF1, p21 in the nuclear and cytoplasmic fractions of indicated lines at Day 0 of Dox/Auxin treatment. Black arrows indicate the endogenous proteins of interest and red arrows indicate the endogenous AID-tagged proteins of interest. (B) Immunoblots of p21, H2AX, γ-H2AX, full-length (FL) PARP and cleaved PARP in the indicated lines at Day 0 of Dox/Auxin treatment. (C) Plots of the master regulator (MR) activity of c-Myc and TFAP4 using the VIPER (Virtual Inference of Protein-activity by Enriched Regulon analysis) algorithm at the indicated time points. (C) Immunoblots of c-Myc and TFAP4 in the indicated conditions.

**Figure S5.**
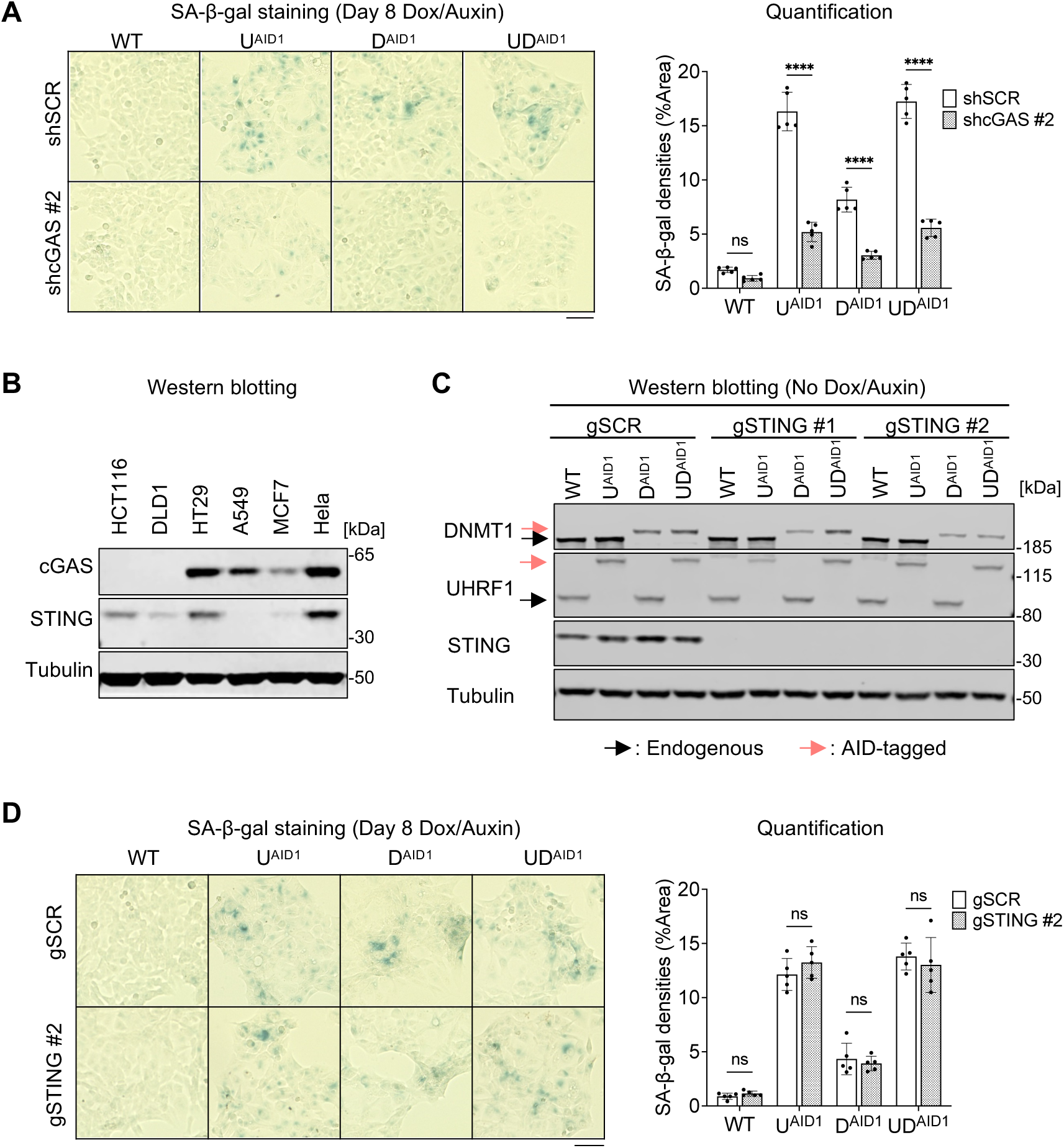
Additional experiments on cGAS and STING in senescence induced by loss of DNA methylation. (A) SA-β-gal staining (left panel) and quantification (right panel) of the indicated lines. N = 5 fields of view. (B) Western blotting of cGAS and STING in various cancer cell lines. (C) Western blot analysis of DNMT1, UHRF1, and STING in the indicated lines untreated with Dox/Auxin. Black arrows indicate the endogenous protein of interest and red arrows indicate the endogenous AID-tagged protein of interest. (D) SA-β-gal staining (left panel) and quantification (right panel) of the indicated lines. N = 5 fields of view. All scale bars are 50 µm. Data of (A) and (D) are presented as mean ± SD and analyzed by two-way ANOVA with Sidak’s multiple comparisons test. We use the following convention: **** p < 0.0001, ns: non-significant.

**Figure S6.**
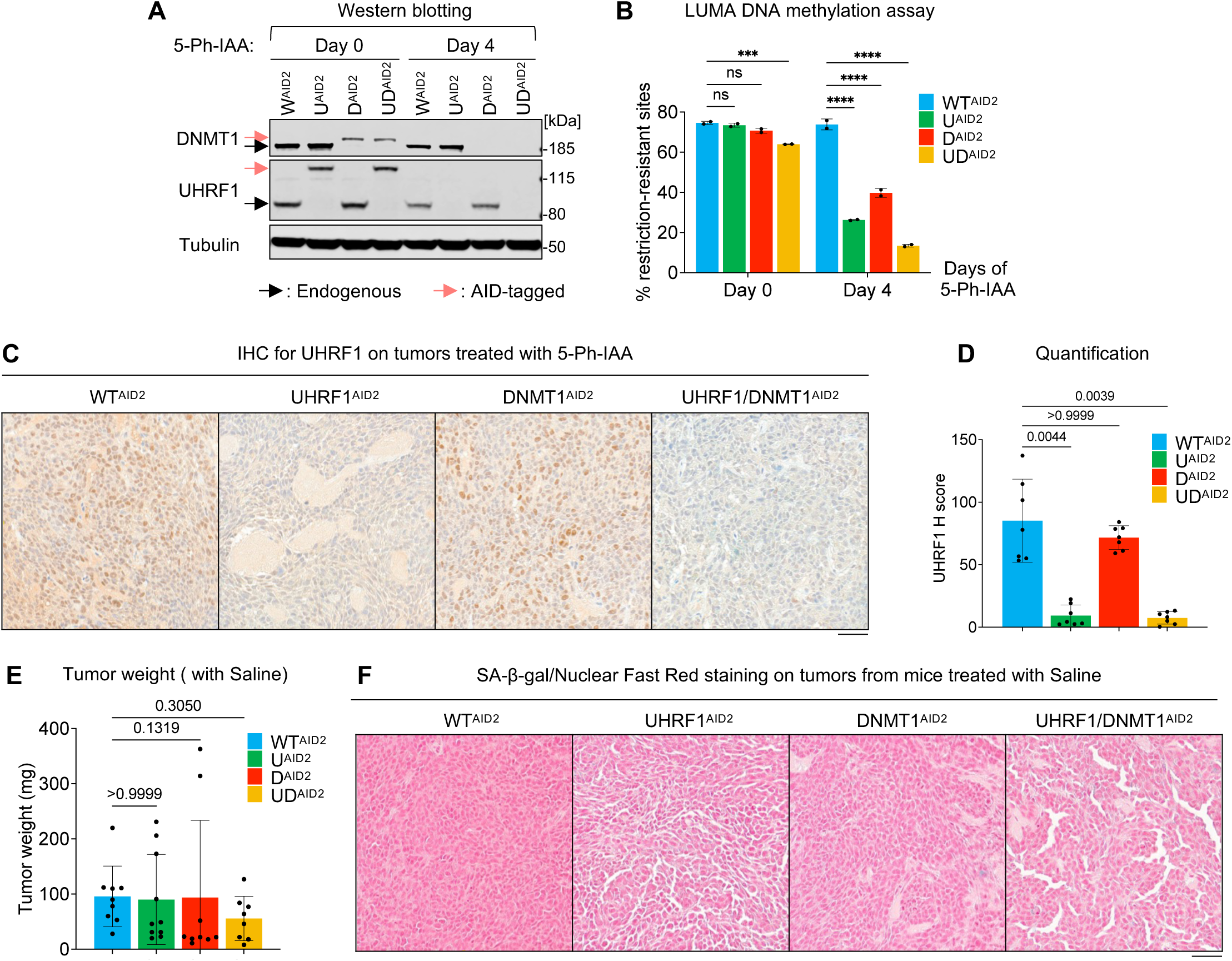
Validation and senescence assessment of HCT116 AID2 system *in vitro* and *in vivo*. (A) Immunoblots of UHRF1 and DNMT1 in HCT116 AID2 lines with and without 5-Ph-IAA treatment. Black arrows indicate the endogenous proteins of interest and red arrows indicate the endogenous AID-tagged proteins of interest. (B) Quantification of the DNA methylation level in each HCT116 AID2 line by LUminometric Methylation Assay (LUMA). N = 2 biological replicates. (C) IHC staining of UHRF1 in indicated conditions. (D) Quantification of UHRF1 IHC using H score. N = 7 fields of view at 400× original magnification. (E) Tumor weight assessment of the mice treated with Saline control. N = 9 tumors for WT^AID2^, 10 tumors for U^ADI2^, 9 tumors for D^AID2^, and 8 tumors for UD^AID2^. (F) Representative images of SA-β-gal staining in indicated control conditions. Nuclear Fast Red was used for counterstaining. All scale bars are 50 µm. All data are presented as mean ± SD. Data of (B) are analyzed by two-way ANOVA and Sidak’s multiple comparisons test. Data of (D) and (E) are analyzed by Kruskal-Wallis test and Dunn’s multiple comparisons test. We use the following convention: *** p < 0.001, **** p < 0.0001, ns: non-significant.

